# An alternative UPF1 isoform drives conditional remodeling of nonsense-mediated mRNA decay

**DOI:** 10.1101/2021.01.27.428318

**Authors:** Sarah E. Fritz, Soumya Ranganathan, Clara D. Wang, J. Robert Hogg

## Abstract

The nonsense-mediated mRNA decay (NMD) pathway monitors translation termination to degrade transcripts with premature stop codons and regulate thousands of human genes. Here we show that an alternative mammalian-specific isoform of the core NMD factor UPF1, termed UPF1_LL_, enables condition-dependent remodeling of NMD specificity. Previous studies indicate that the extension of a conserved regulatory loop in the UPF1_LL_ helicase core confers a decreased propensity to dissociate from RNA upon ATP hydrolysis relative to the major UPF1 isoform, designated UPF1 _SL_. Using biochemical and transcriptome-wide approaches, we find that UPF1_LL_ overcomes the protective RNA binding proteins PTBP1 and hnRNP L to preferentially bind and down-regulate long 3’UTRs normally shielded from NMD. Unexpectedly, UPF1_LL_ supports induction of NMD on new populations of substrate mRNAs in response to activation of the integrated stress response and impaired translation efficiency. Thus, while canonical NMD is abolished by moderate translational repression, UPF1_LL_ activity is enhanced, providing a mechanism to rapidly rewire NMD specificity in response to cellular stress.

## Introduction

Nonsense-mediated mRNA decay (NMD) is an evolutionarily conserved mRNA quality-control pathway that degrades transcripts undergoing premature translation termination (Lavysh & Neu-Yilik, 2020; Smith & Baker, 2015). In addition, NMD performs a regulatory role by governing the turnover of ~5-10% of the transcriptome, including mRNAs with upstream open reading frames, introns downstream of the stop codon, or long 3’ untranslated regions (UTRs) (Kishor *et al*, 2019). Despite extensive studies of the large impact of NMD on the transcriptome, the mechanisms by which the pathway selects its regulatory targets are poorly understood.

A suite of conserved NMD factors acts in concert with general mRNA binding proteins and RNA decay enzymes to identify and degrade target mRNAs. The RNA helicase UPF1 is a central coordinator of the NMD pathway, as it directly binds mRNA and functions at multiple steps in the selection and degradation of target transcripts (Kim & Maquat, 2019). Additional core NMD factors UPF2 and UPF3 promote UPF1 activity and link UPF1 to the exon junction complex (EJC), which strongly stimulates decay (Chamieh *et al*, 2008; Le Hir *et al*, 2000, 2000a). In many eukaryotes, NMD execution also depends on the SMG1, 5, 6, and 7 proteins (Page *et al*, 1999; Pulak & Anderson, 1993; Hodgkin *et al*, 1989; Causier *et al*, 2017). Phosphorylation of UPF1 by the SMG1 kinase strongly promotes decay (Kashima *et al*, 2006), as phosphorylated UPF1 recruits the SMG6 endonuclease and/or general decapping and deadenylation enzymes through the SMG5/7 heterodimer (Eberle *et al*, 2009; Huntzinger *et al*, 2008; Loh *et al*, 2013).

In addition to the functions of specialized NMD proteins, substrate selection and degradation by the NMD pathway requires the translation termination machinery to detect in-frame stop codons (Karousis & Mühlemann, 2018). Although the exact molecular details remain to be elucidated, widely accepted models of NMD state that interactions between core NMD factors and a terminating ribosome are necessary for decay. Because of the strict dependence of NMD on translation termination, decay efficiency of canonical NMD targets is expected to be tightly linked to translation efficiency. However, in contrast to this prevailing model, there is evidence that NMD efficiency for some targets is actually enhanced during conditions of impaired translation (Martinez-Nunez *et al*, 2017). These data warrant a more extensive investigation into the role of translation in shaping target specificity by the NMD pathway, particularly during changing physiological conditions.

The ability of UPF1 to bind and hydrolyze ATP is critical for the selection and degradation of potential NMD substrates (Lee *et al*, 2015; Franks *et al*, 2010). Numerous studies have provided evidence that the affinity of UPF1 for RNA is reduced by ATP binding and hydrolysis, in a manner dependent on an 11 amino acid regulatory loop in domain 1B of the helicase core that protrudes into the RNA binding channel (Cheng *et al*, 2007; Chakrabarti *et al*, 2011; Gowravaram *et al*, 2018; Chamieh *et al*, 2008; Fiorini *et al*, 2013; Czaplinski *et al*, 1995; Weng *et al*, 1998). Intriguingly, mammals undergo an alternative splicing event to express two UPF1 isoforms that differ only in length of the regulatory loop (Fig 1A). Almost all NMD studies to date have focused on the more abundant UPF1 “short loop” isoform (designated herein UPF1_SL_), which contains the 11 amino acid regulatory loop that most potently weakens the affinity of UPF1 for RNA in the presence of ATP. Alternative 5’ splice site usage in exon 7 of UPF1 generates a second UPF1 isoform that extends the regulatory loop to 22 amino acids. This naturally occurring UPF1 “long loop” isoform (designated herein UPF1_LL_), which represents ~15-25% of total UPF1 mRNA in diverse cell and tissue types (Fig 1B), has increased catalytic activity and a higher affinity for RNA in the presence of ATP than the UPF1_SL_ isoform (Gowravaram *et al*, 2018). It is unknown whether the differential biochemical properties of the UPF1_LL_ isoform affect NMD specificity in cells.

**Figure 1.**
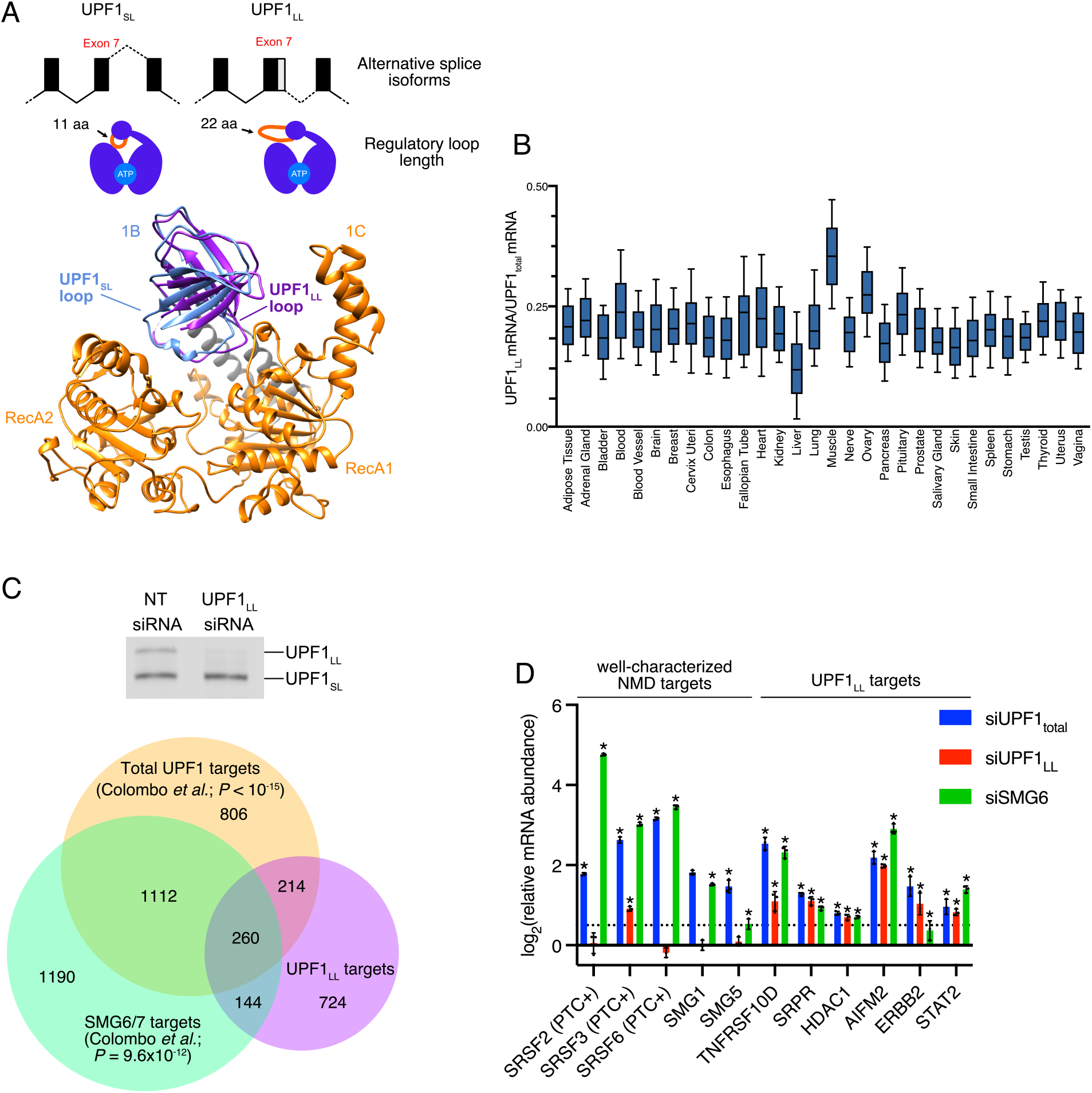
Alternative UPF1_LL_ splice isoform contributes to NMD under normal cellular conditions. **A. (Top)** Schematic representation of alternative 5’ splice site usage in exon 7 of mammalian UPF1 that results in two UPF1 protein isoforms that differ in length of the regulatory loop within the helicase core. **(Bottom)** Ribbon diagram of human UPF1_SL_ helicase core overlaid with that of human UPF1_LL_. The regulatory loop in domain 1B is indicated for each UPF1 variant, with light blue representing the 11 amino acid regulatory loop of human UPF1_SL_ and purple indicating the 22 amino acid regulatory loop of human UPF1_LL_. This figure was generated using Protein Data Bank accessions 2XZP and 6EJ5 (Chakrabarti et al., 2011; Gowravaram et al., 2018). **B.** Box plot of the fraction of UPF1_LL_ /UPF1_total_ mRNA expressed in indicated human tissues, as determined from the Genotype-Tissue Expression (GTEx) project. Boxes indicate interquartile ranges, and bars indicate 10-90% ranges. **C. (Top)** Semiquantitative RT-PCR of UPF1_SL_ or UPF1_LL_ transcript levels following transfection of HEK-293 cells with a non-targeting (NT) siRNA or a siRNA that specifically targets the UPF1_LL_ isoform. **(Bottom)** Venn diagram (to scale) of overlapping targets identified from RNA-seq following UPF1_LL_ knockdown (this dataset), total UPF1 knockdown, or SMG6/7 double knockdown and rescue (Colombo et al., 2017). Depicted are genes that increased in abundance at least 1.4-fold (FDR < 0.05) with UPF1_LL_-specific knockdown and their overlap with genes that increased in abundance (FDR < 0.05) with total UPF1 knockdown or genes that increased in abundance with SMG6/7 double knockdown and were significantly rescued by expression of SMG6 or SMG7 (SMG6/7 targets). *P* values indicate enrichment of genes that increased in abundance at least 1.4-fold (FDR < 0.05) with UPF1_LL_-specific knockdown among those regulated by total UPF1 and SMG6/7, as determined by Fisher’s exact test. **D.** RT-qPCR analysis of indicated transcripts following transfection of HEK-293 cells with siRNAs that target both UPF1 isoforms (UPF1_total_), the UPF1_LL_-specific isoform, or the NMD endonuclease SMG6. Relative fold changes are in reference to non-targeting siRNA. Significance of total UPF1, UPF1_LL_’ or SMG6 knockdown was compared to the NT siRNA control. Asterisk (*) indicates *P* < 0.0001, as determined by two-way ANOVA. Error bars indicate SD (n = 3). Dashed line indicates log_2_(fold change) of +0.5. PTC+ indicates the use of primers specific to transcript isoforms with validated poison exons (Lareau et al., 2007; Ni et al., 2007). See also Table S2 for Pvalues associated with each statistical comparison.

Here, we show that the UPF1_LL_ isoform gives the mammalian NMD pathway the latent ability to remodel NMD target specificity in response to changing physiological conditions. We identify that UPF1_LL_ can overcome inhibition by polypyrimidine tract binding protein 1 (PTBP1) and heterogeneous nuclear ribonucleoprotein L (hnRNP L) to preferentially associate with and down-regulate long 3’UTRs normally shielded from NMD. Unexpectedly, we find that UPF1_LL_ activity is enhanced in conditions of reduced translation efficiency, including during the integrated stress response. mRNAs subject to UPF1_LL_-dependent down-regulation upon translation inhibition include hundreds of mRNAs not normally targeted by NMD, many of which are protected by PTBP1 and hnRNP L. Together our data support that human cells use the UPF1_LL_ isoform to conditionally alter which mRNAs are selected and degraded by the NMD pathway, expanding the scope of NMD in mammalian gene expression control.

## Results

### UPF1_LL_ contributes to NMD under normal cellular conditions

To specifically interrogate the cellular functions of the UPF1_LL_ isoform, we developed an siRNA that efficiently down-regulates UPF1_LL_ mRNA without perturbing the expression of the major UPF1_SL_ isoform (Fig 1C (top) and S1A). As an initial analysis of UPF1_LL_ functions, we treated human HEK-293 cells with the UPF1_LL_-specific siRNA and performed total RNA-seq. Differential expression analysis identified 1621 genes that were at least 1.4-fold more highly expressed upon UPF1_LL_ knockdown, out of a total population of 13,668 genes analyzed, indicating a role for endogenous UPF1_LL_ in gene expression regulation (for complete characteristics of the RNA-seq dataset, see Table S1).

To ask whether these changes in gene expression were due to activity of UPF1_LL_ in the NMD pathway, we compared our UPF1_LL_-knockdown RNA-seq dataset with a published catalog of high-confidence NMD targets (Colombo *et al*, 2017). We observed significant overlaps among the population of genes induced by UPF1_LL_ depletion and those previously determined to be repressed by UPF1, SMG6, or SMG7 (Fig 1C, bottom), with 618 of the 1621 putative UPF1_LL_ targets represented in the published NMD target catalog. These results strongly implicate endogenous UPF1_LL_ in the overall activities of NMD under normal cellular conditions and are corroborated by quantitative RT-PCR experiments showing up-regulation of select transcripts upon UPF1_LL_, total UPF1, or SMG6 knockdown (Fig 1D).

Specific analysis of well-characterized NMD targets revealed that UPF1_LL_ knockdown, unlike total UPF1 and SMG6 knockdown, did not affect the levels of SMG1 and SMG5 mRNAs, both of which have been shown by several groups to undergo NMD due to their unspliced long 3’UTRs (Singh *et al*, 2008; Huang *et al*, 2011; Yepiskoposyan *et al*, 2011; Hogg & Goff, 2010). To assess the role of UPF1_LL_ in regulating EJC-stimulated NMD, we examined a set of 1640 transcript isoforms predicted to contain stop codons at least 50 nt upstream of the final exon-exon junction (premature termination codons; PTCs), using transcripts derived from the same genes but predicted to contain normal termination codons as controls. Depletion of total UPF1 levels caused elevated expression of PTC-containing transcript isoforms relative to control PTC-free isoforms (Fig S1B). In contrast, selective depletion of UPF1_LL_ did not systematically affect the abundance of mRNAs predicted to contain PTCs. Consistent with these transcriptome-wide observations, knockdown of UPF1_LL_ had no effect on the levels of well-characterized PTC-containing SRSF2 and SRSF6 transcripts, and increased SRSF3 PTC transcript levels to a much smaller extent (~1.9-fold) than total UPF1 (~6.2-fold) or SMG6 (~8.1-fold) knockdown (Fig 1D; (Ni *et al*, 2007; Lareau *et al*, 2007). Based on these data, we conclude that UPF1_LL_ is dispensable for the turnover of several well-characterized EJC- and 3’UTR-stimulated targets but is uniquely required for a subset of cellular UPF1 activities.

### Enhanced UPF1_LL_ binding to NMD-resistant transcripts

The observation that specific depletion of UPF1_LL_ affected a select subpopulation of NMD targets indicated it has distinct cellular functions from those of the major UPF1_SL_ isoform. To gain insight into how the biochemical activities of the two UPF1 isoforms differ, we performed affinity purification followed by RNA-seq (RIP-seq) of each UPF1 variant (Fig 2A).

**Figure 2.**
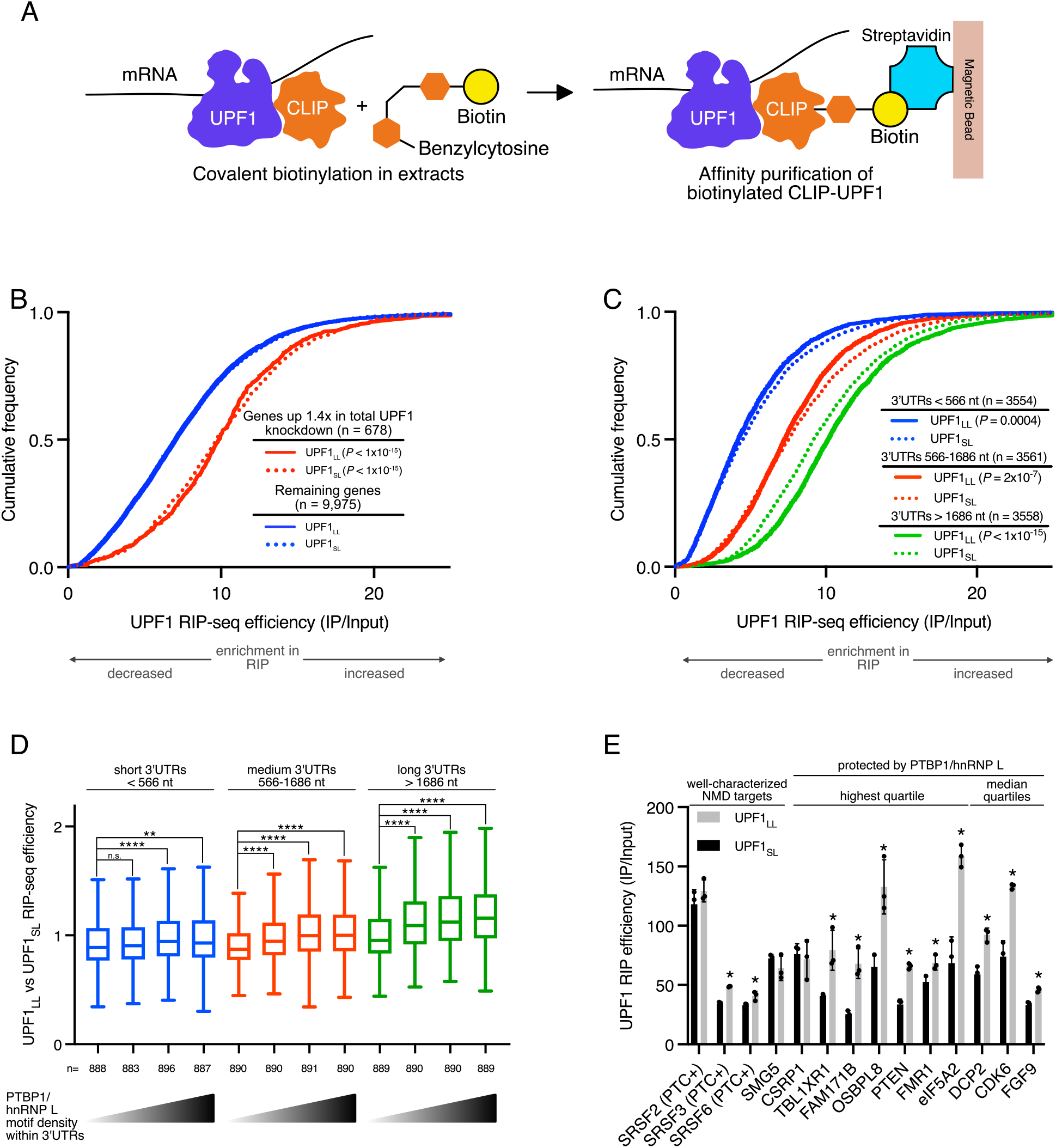
UPF1_LL_ can bind NMD-protected transcripts. **A.** Scheme for the CLIP-UPF1 affinity purification (RIP) assay. **B.** CDF plot of recovered mRNAs as determined from RIP-seq efficiency in CLIP-UPF1_LL_ or CLIP- UPF1_SL_ affinity purifications. mRNAs were binned by sensitivity to total UPF1 knockdown (Baird et al., 2018). Statistical significance was determined by K-S test. **C.** CDF plot of recovered mRNAs as determined from RIP-seq efficiency in CLIP-UPF1_LL_ or CLIP- UPF1_SL_ affinity purifications. mRNAs were binned according to 3’UTR length. Significance of differential recovery in CLIP-UPF1_LL_ RIP was determined by K-S test of comparison to that in CLIP-UPF1_SL_. **D.** Box plot of recovered mRNAs, as determined from RIP-seq efficiency in CLIP-UPF1_LL_ vs CLIP-UPF1_SL_ affinity purifications. mRNAs were binned according to 3’UTR length and then equally subdivided by PTBP1 and/or hnRNP L motif density within the 3’UTR, as indicated by the gradient triangles. Statistical significance was determined by K-W test, with Dunn’s correction for multiple comparisons (** *P* < 0.002, **** *P* < 5×10^−6^). Boxes indicate interquartile ranges, and bars indicate Tukey whiskers. **E.** RT-qPCR analysis of indicated transcripts from UPF1 RIP-seq experiments. Relative fold enrichment was determined by dividing the recovered mRNA by its corresponding input amount. Significance of differential recovery in CLIP-UPF1_LL_ RIP was determined by comparison to that in CLIP-UPF1_SL_. Asterisk (*) indicates *P* < 0.05, as determined by unpaired Student’s t-test. Error bars indicate SD (n = 3). For protected mRNAs, the PTBP1/hnRNP L motif density bin of the 3’UTR is indicated. PTC+ indicates the use of primers specific to transcript isoforms with validated poison exons (Lareau et al., 2007; Ni et al., 2007). See also Table S2 for Pvalues associated with each statistical comparison.

For these studies, we engineered HEK-293 stable cell lines to express CLIP-tagged UPF1_LL_ or UPF1_SL_, with a GFP-expressing stable line as a control. We elected to express CLIP-tagged UPF1 constructs, as the CLIP tag can be used to covalently biotinylate tagged proteins of interest (Gautier *et al*, 2008) for efficient isolation by streptavidin affinity purification. We have previously used this system to show that biotinylated CLIP-UPF1_SL_ isolated from human cells preferentially associates with NMD-susceptible mRNA isoforms (Kishor *et al*, 2020).

CLIP-UPF1 complexes were isolated from whole cell extracts by streptavidin affinity purification using a CLIP-biotin substrate, with GFP-expressing cell lines as a negative control for interaction specificity (Fig S2A). Bound RNAs were then extracted and used for sequencing library preparation. Because recovery of RNA from GFP samples was at least 100-fold lower than from CLIP-UPF1 affinity purifications (Table S3), only UPF1 samples were analyzed by RNA-seq. UPF1 occupancy was assessed by normalizing the abundance of transcripts in RIP-seq samples to their abundance in total RNA-seq (hereafter referred to as UPF1 RIP-seq efficiency).

The two UPF1 isoforms both strongly enriched mRNAs up-regulated by total UPF1 depletion (Fig 2B), consistent with previous observations of preferential UPF1 association with decay targets (Hurt *et al*, 2013; Zünd *et al*, 2013; Kishor *et al*, 2018, 2020). UPF1 has been shown to specifically accumulate on 3’UTRs due to its active displacement from coding regions by translating ribosomes and its nonspecific RNA binding activity (Hogg & Goff, 2010; Hurt *et al*, 2013; Baker & Hogg, 2017; Kurosaki *et al*, 2014; Zünd *et al*, 2013). Subdivision of the transcriptome according to 3’UTR lengths (first tertile: < 566 nt; second tertile: 566-1686 nt; third tertile: > 1686 nt) revealed that the efficiency of mRNA co-purification with both UPF1_SL_ and UPF1_LL_ increased with 3’UTR length (Fig 2C), reinforcing the conclusion that UPF1 binding correlates with 3’UTR length. Distinctly, however, CLIP-UPF1_LL_ more efficiently recovered the longest class of 3’UTRs than UPF1_SL_.

Transcripts with long 3’UTRs represent a large population of potential NMD targets (Hurt *et al*, 2013; Yepiskoposyan *et al*, 2011), only some of which are degraded by the pathway under normal conditions (Toma *et al*, 2015). Providing a biochemical mechanism to explain evasion of long 3’UTRs from decay, we have identified hundreds to thousands of mRNAs shielded by the protective RNA-binding proteins (RBPs) PTBP1 and hnRNP L Ge *et al*, 2016; Kishor *et al*, 2018, 2020). In our previous work, we showed that increased PTBP1 and/or hnRNP L motif binding density within the 3’UTR correlates with reduced UPF1_SL_ binding and recovery of mRNAs in UPF1_SL_ RIP-seq studies (Ge *et al*, 2016; Kishor *et al*, 2018; Fritz *et al*, 2020). The observation that UPF1_LL_ more efficiently recovers the longest class of 3’UTRs led us to ask whether mRNAs protected by PTBP1 and/or hnRNP L are differentially associated with UPF1_LL_ versus UPF1_SL_. Subdivision of the transcriptome first by 3’UTR length and then according to the density of PTBP1 and hnRNP L binding sites within the 3’UTR revealed that transcripts with long 3’UTRs and moderate or high densities of protective protein binding sites were more efficiently recovered by CLIP-UPF1_LL_ than CLIP-UPF1_SL_ (Fig 2D). This preferential recovery of long 3’UTRs with moderate or high densities of protective protein binding by CLIP-UPF1_LL_ was similarly observed when PTBP1 and/or hnRNP L motif densities were restricted to the first 400 nt of the 3’UTR (Fig S2B), which we previously established as a strong feature driving protection and reduced UPF1_SL_ binding (Ge *et al*, 2016; Kishor *et al*, 2018). Quantitative RT-PCR of select transcripts confirmed these transcriptome-wide RIP-seq results (Fig 2E). Together, our findings indicate that the distinct biochemical properties of UPF1_LL_ allow it to circumvent the protective effects of PTBP1 and/or hnRNP L to associate with otherwise protected mRNAs.

### UPF1_LL_ is less sensitive to PTBP1-mediated inhibition of translocation

We have proposed that the protective RBPs PTBP1 and hnRNP L exploit the tendency of UPF1 to release RNA upon ATP binding and hydrolysis to promote UPF1 dissociation from potential NMD substrates prior to decay induction (Fritz *et al*, 2020). In support of this model, deletion of the regulatory loop, which mediates ATPase-dependent dissociation, rendered UPF1_SL_ less sensitive to PTBP1 inhibition *in vitro* (Fritz *et al*, 2020). Importantly, both the physiological UPF1_LL_ isoform and the engineered UPF1 variant containing a regulatory loop deletion exhibit a greater affinity for RNA in the presence of ATP than the PTBP1-sensitive UPF1_SL_ isoform (Gowravaram *et al*, 2018). We therefore hypothesized that UPF1_LL_ can mimic the ability of the loop truncation mutant to overcome negative regulation by PTBP1.

We recently established a real-time assay to monitor UPF1 translocation activity (Fritz *et al*, 2020). In this assay, UPF1 translocation and duplex unwinding causes a fluorescently labeled oligonucleotide to be displaced from the assay substrate (Fig 3A, left). An excess of complementary oligonucleotide labeled with a dark quencher is provided in the reaction, causing a decrease in fluorescence with increased displacement of the labeled oligonucleotide by UPF1. Inhibition of UPF1 translocation results in sustained fluorescence over time, allowing for the determination of inhibitory effects of PTBP1 on UPF1 unwinding activity. Using this assay in our previous work, we showed PTBP1 impairs UPF1_SL_ unwinding activity in a manner consistent with a mechanism in which PTBP1 inhibits UPF1 translocation rather than initial binding (Fritz *et al*, 2020). This inhibitory effect on UPF1 translocation activity was specific to PTBP1 and was not observed in the presence of the high-affinity RNA binding *Pseudomonas* phage 7 coat protein, supporting the conclusions that the protective proteins specifically promote the dissociation of UPF1 and that our assay can robustly assess inhibitors of UPF1 unwinding activity.

**Figure 3.**
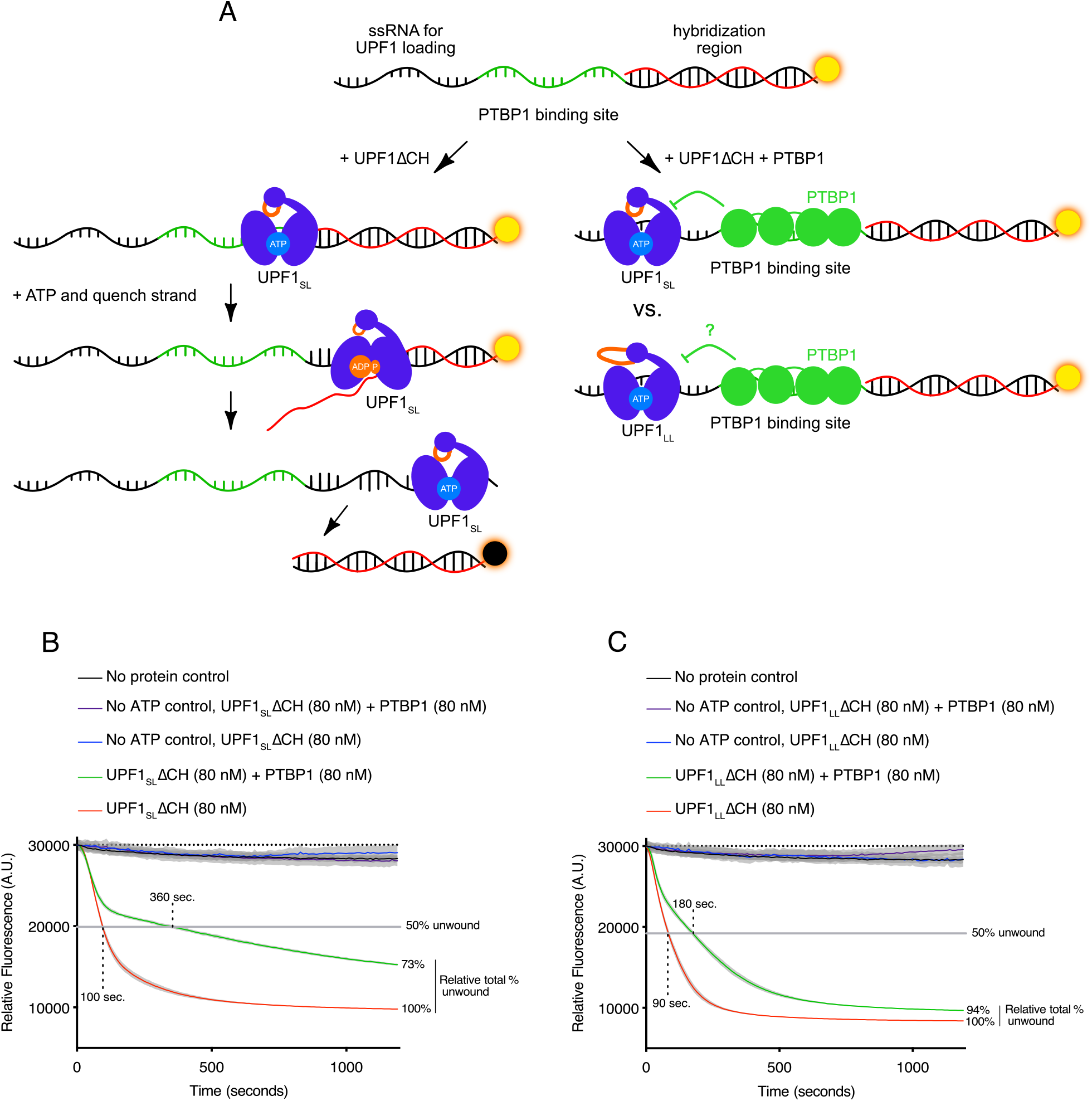
UPF1_LL_ overcomes the translocation inhibition by PTBP1. **A.** Scheme of the fluorescence-based unwinding assay to monitor UPF1 translocation in real time (Fritz et al., 2020). An RNA substrate harboring a high-affinity PTBP1 binding site is incubated with highly purified UPF1ΔCH in the absence and presence of equal molar amounts of highly purified PTBP1. Upon the addition of ATP, UPF1 translocation results in a decrease in fluorescence due to displacement of a labeled, duplexed oligonucleotide and subsequent quenching by a trap strand. **B.** UPF1_SL_ΔCH translocation along an RNA substrate harboring a high-affinity PTBP1 binding site in the absence and presence of PTBP1. Time to 50% unwound and relative total % unwound at end-of-assay (1200 sec) are indicated. Results of four technical replicates are shown for each dataset and represent at least three independent experiments. Shaded region indicates SD. **C.** Results as in (B) but with UPF1_LL_ΔCH.

We therefore leveraged this system to compare UPF1_LL_ versus UPF1_SL_ translocation on a duplexed RNA substrate harboring a high affinity PTBP1 binding site (Fig 3A, right) (Fritz *et al*, 2020). For these experiments, we compared the activity of highly purified UPF1 proteins containing the helicase core but lacking the autoinhibitory N-terminal cysteine-histidine domain (UPF1ΔCH) (Fig S3A and B) (Chakrabarti *et al*, 2011; Fiorini *et al*, 2012; Fritz *et al*, 2020). UPF1_SL_ΔCH exhibited robust unwinding activity in the absence of PTBP1, displacing 50% of the duplexed oligonucleotide in 100 seconds (Fig 3B). This translocation activity of UPF1_SL_ΔCH was dependent upon the addition of ATP, as previously demonstrated (Fritz *et al*, 2020). Addition of PTBP1 substantially impaired UPF1_SL_ΔCH unwinding activity, reducing both the rate at which the oligonucleotide was displaced (requiring 360 seconds to attain the half-maximal unwinding value reached by UPF1_SL_ΔCH alone) and the overall extent of unwinding (73% of the UPF1_SL_ΔCH total at the end of the assay).

UPF1_LL_ΔCH also exhibited robust unwinding activity in the absence of PTBP1, displacing 50% of the duplexed oligonucleotide in 90 seconds in an ATP-dependent manner (Fig 3C). The observed enhancement in UPF1_LL_ΔCH translocation activity over UPF1_SL_ΔCH is consistent with previous reports of increased catalytic activity of the UPF1_LL_ isoform relative to UPF1_SL_ (Gowravaram *et al*, 2018). In contrast to UPF1_SL_ΔCH, UPF1_LL_ΔCH maintained robust unwinding activity in the presence of PTBP1, displacing 50% of the duplexed oligonucleotide by 180 seconds and achieving 94% total duplex unwinding at the end of the assay. These results indicate that UPF1_LL_ can overcome the translocation inhibition by PTBP1, reinforcing the conclusion that PTBP1-mediated UPF1 inhibition depends on the clash between the UPF1 regulatory loop and RNA.

### UPF1_LL_ overexpression down-regulates mRNAs normally protected from NMD

Having shown that UPF1_LL_ exhibits enhanced association with normally protected mRNAs in cells (Fig 2) and is able to overcome inhibition by PTBP1 *in vitro* (Fig 3), we next asked whether the distinct biochemical properties of UPF1_LL_ allow it to promote the degradation of mRNAs that normally evade UPF1-dependent decay. Because endogenous UPF1_LL_ mRNA is expressed at ~15-25% of total UPF1 mRNA levels, we hypothesized that the cellular activities of UPF1_LL_ are likely constrained by low basal expression. To test this hypothesis, we used the CLIP-UPF1 stable lines generated for the RIP-seq studies (Fig 2), which inducibly overexpress CLIP-UPF1_SL_ or CLIP-UPF1_LL_ to levels ~5x greater than that of the total UPF1 pool (Fig S4A). Analysis of well-characterized NMD substrate levels following knockdown of total endogenous UPF1 and rescue with the siRNA-resistant CLIP-UPF1 constructs confirmed that both CLIP-tagged UPF1 isoforms were equally able to function in NMD (Fig S4B).

Having established that the CLIP-UPF1 stable lines could be used to assess potential gain-of-function activities with UPF1 overexpression, we analyzed total RNA-seq from the CLIP-UPF1 and GFP-expressing control cells. Because UPF1_LL_ showed a greater propensity to recover the longest class of 3’UTRs than UPF1_SL_ (Fig 2C), we first evaluated the effects of UPF1_SL_ and UPF1_LL_ overexpression by subdividing the transcriptome according to 3’UTR lengths (short: < 566 nt; medium: 566-1686 nt; long: > 1686 nt). Overexpression of UPF1_SL_ led to mRNA abundance changes that were tightly distributed around zero for all 3’UTR length classes (10-90% log_2_FC ranges: short, −0.18 to 0.23; medium, −0.19 to 0.21; long, −0.18 to 0.18). In contrast, overexpression of UPF1_LL_ induced greater variability in gene expression for all 3’UTR length classes (10-90% log2FC ranges: short, −0.27 to 0.38; medium, −0.29 to 0.37; long, −0.44 to 0.29). Moreover, consistent with the RIP-seq findings, overexpression of UPF1_LL_ preferentially promoted the downregulation of mRNAs with the longest 3’UTRs (Fig 4A). Further subdivision of long 3’UTRs according to their density of PTBP1 and/or hnRNP L binding sites revealed that mRNAs with long 3’UTRs and a moderate or high density of binding sites for the protective proteins were significantly down-regulated by UPF1_LL_ overexpression relative to those with a low density of binding sites (Fig 4B). In contrast, overexpression of UPF1_SL_ did not systematically perturb expression of 3’UTRs of any motif density class. These results support the hypothesis that UPF1_LL_ is biochemically capable of overcoming inhibition by PTBP1 and hnRNP L, but that the cellular activities of UPF1_LL_, unlike those of UPF1_SL_, are limited by low expression levels.

**Figure 4.**
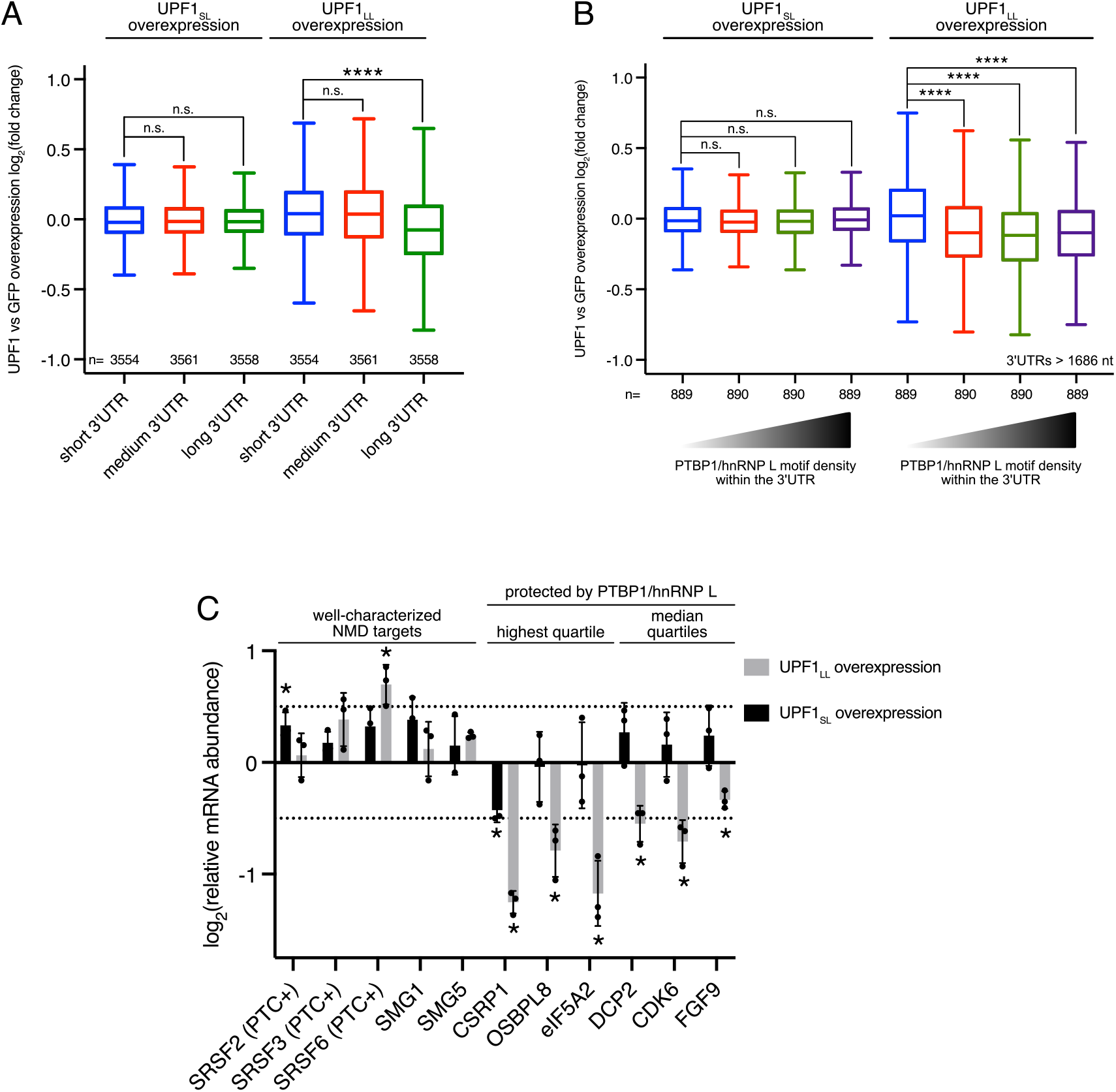
UPF1_LL_ can down-regulate mRNAs normally protected from NMD. **A.** Box plot of relative mRNA abundance as determined from RNA-seq following CLIP-UPF1_SL_ or CLIP- UPF1_LL_ overexpression. mRNAs were binned by 3’UTR length (short < 566 nt; medium 566-1686 nt; long > 1686 nt). Statistical significance was determined by K-W test, with Dunn’s correction for multiple comparisons (**** *P* < 1×10-**15).** Boxes indicate interquartile ranges, and bars indicate Tukey whiskers. **B.** Box plot of relative mRNA abundance as determined from RNA-seq following CLIP-UPF1_SL_ or CLIP- UPF1_LL_ overexpression. mRNAs with long 3’UTRs (> 1686 nt) were equally subdivided by PTBP1 and/or hnRNP L motif density within the 3’UTR, as indicated by the gradient triangles. Statistical significance was determined by K-W test, with Dunn’s correction for multiple comparisons (**** *P* < 1×10^−15^). Boxes indicate interquartile ranges, and bars indicate Tukey whiskers. **C.** RT-qPCR analysis of indicated transcripts from CLIP-UPF1 overexpression RNA-seq experiments. Relative fold changes are in reference to the GFP expressing control line. Significance of CLIP-UPF1_SL_ or CLIP-UPF1_LL_ overexpression was compared to the GFP expressing control line. Asterisk (*) indicates *P* < 0.05, as determined by two-way ANOVA. Error bars indicate SD (n = 3). Dashed lines indicate log_2_(fold change) of ±0.5. For protected mRNAs, the PTBP1/hnRNP L 3’UTR motif density bin is indicated. PTC+ indicates the use of primers specific to transcript isoforms with validated poison exons (Lareau et al., 2007; Ni et al., 2007). See also Table S2 for *P* values associated with each statistical comparison.

To investigate whether the observed changes in mRNA abundance with UPF1_LL_ overexpression were due to enhanced decay, we used REMBRANDTS software to assess changes in mRNA stability based upon differences in the relative abundance of exonic and intronic reads from each gene (Alkallas *et al*, 2017). The effects of UPF1_SL_ overexpression on mRNA abundance were not correlated with changes to mRNA stability (Spearman’s ρ = −0.45; *P* = 10^−5^; Fig S4C), corroborating the previous observation that moderate UPF1_SL_ overexpression does not enhance NMD activity (Huang *et al*, 2011). In contrast, the effects of UPF1_LL_ (Spearman’s ρ = 0.65; *P* < 10^−15^; Fig S4D), with transcripts having long 3’UTRs and a high density of PTBP1 and/or hnRNP L binding motifs showing preferential destabilization with UPF1_LL_ overexpression (Fig S4E and F).

We further corroborated the transcriptome-wide results obtained using RNA-seq by performing RT-qPCR on select transcripts (Fig 4C). Notably, validated mRNAs down-regulated by UPF1_LL_ overexpression include CSRP1, which we have previously established as a long 3’UTR-containing mRNA that undergoes decay upon hnRNP L knockdown or mutation of hnRNP L binding sites in its 3’UTR (Kishor *et al*, 2018). Together, these data support the conclusion that the UPF1_LL_ isoform is able to overcome the protective proteins to promote decay of mRNAs normally shielded from NMD.

### Coordinated downregulation of UPF1_LL_ targets during ER stress and ISR induction

Our *in vitro*, RIP-seq, and overexpression studies suggested that UPF1_LL_ has the biochemical capacity to expand the scope of UPF1-dependent regulation. Based on these observations, we next investigated whether specific physiological conditions might promote changes in NMD target susceptibility by harnessing endogenous UPF1_LL_ activity. Because genes in functionally related pathways are often coordinately regulated at the posttranscriptional level (Keene, 2007), we performed a gene ontology (GO) enrichment analysis (Eden *et al*, 2009) to identify commonalities among UPF1_LL_ targets. We reasoned that functional similarities among the 1621 genes up-regulated in response to UPF1_LL_-specific depletion under normal cellular conditions (Fig 1; hereafter referred to as constitutive UPF1_LL_ targets) might indicate physiological conditions that influence endogenous UPF1_LL_ activity. This analysis revealed a high degree of enrichment among constitutive UPF1_LL_ targets for genes encoding proteins that rely on the endoplasmic reticulum (ER) for biogenesis. In total, 768 of the 1621 genes up-regulated by UPF1_LL_ depletion are annotated by UniProt as encoding integral membrane, secreted, and/or signal peptide-containing proteins (Fig 5A and Table S4). We also used a previous survey of ER-localized translation (Jan *et al*, 2014) to corroborate the results of the GO analysis, finding that many UPF1_LL_ target mRNAs were indeed found to be preferentially translated at the ER (Fig S5A).

**Figure 5.**
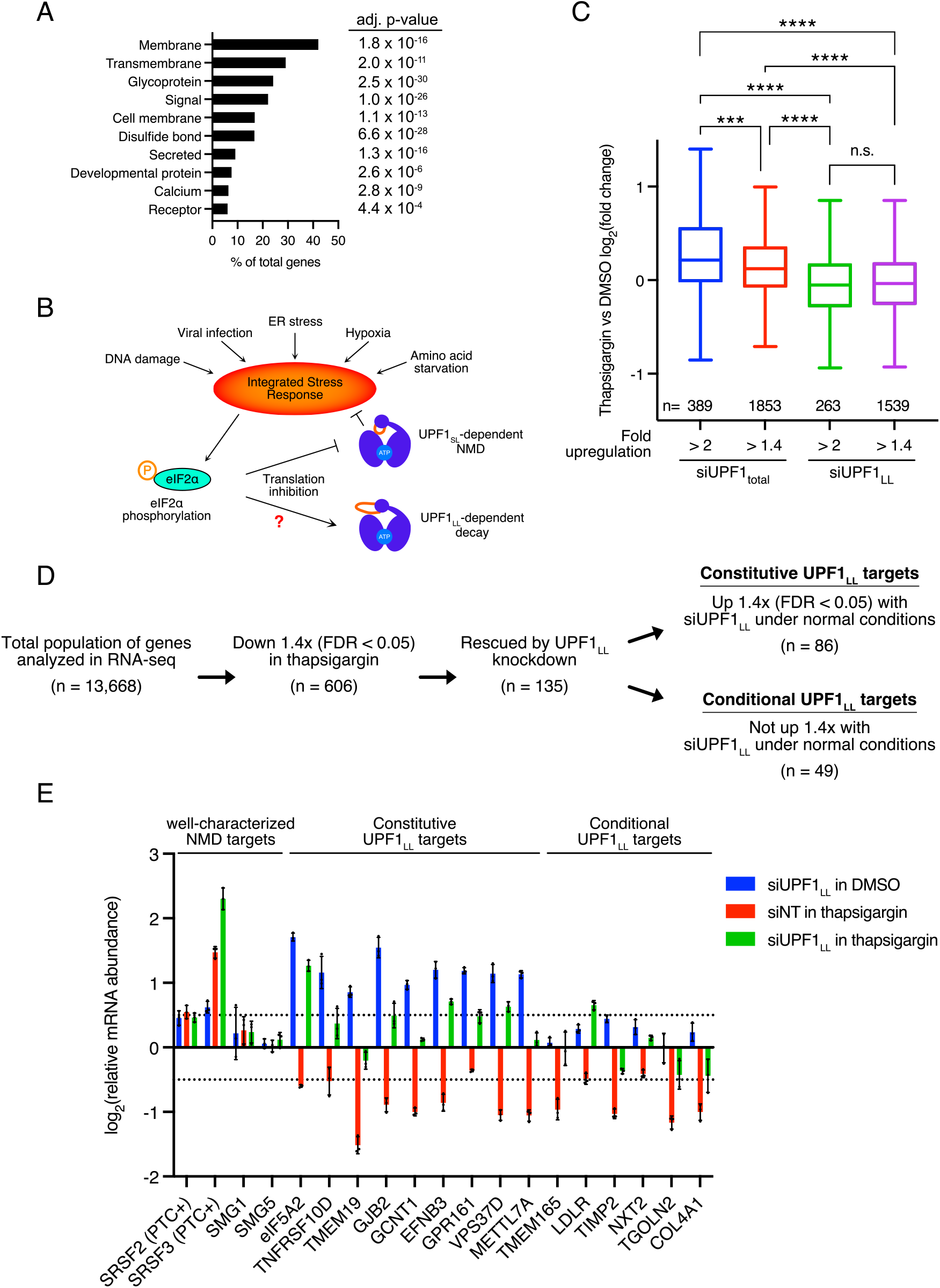
UPF1_LL_ conditionally remodels NMD target selection during ER stress and induction of the ISR. **A.** Gene ontology analysis of 1621 genes that increased in expression at least 1.4-fold upon UPF1_LL_ - specific knockdown in HEK-293 cells under normal cellular conditions. Genes may map to multiple categories. **B.** Scheme for activation of the integrated stress response (ISR) and effects on UPF1-dependent decay. **C.** Box plot of relative mRNA abundance as determined from RNA-seq following treatment of HEK-293 cells with 1 *μ*M thapsigargin for 6hr. mRNAs were binned according to sensitivity to total UPF1 (UPF1_total_) or UPF1_LL_-specific knockdown under basal conditions. Statistical significance was determined by K-W test, with Dunn’s correction for multiple comparisons (*** *P* = 0.0001; **** *P* < 6×10^−14^). Boxes indicate interquartile ranges, and bars indicate Tukey whiskers. **D.** RNA-seq analysis of HEK-293 cells identifies populations of genes that decreased in abundance with thapsigargin treatment and were rescued by UPF1_LL_-specific knockdown. ndicated are genes that increased in abundance at least 1.4-fold (FDR < 0.05) with UPF1_LL_-specific knockdown under normal conditions (Constitutive UPF1_LL_ targets) and those that did not (Conditional UPF1_LL_ targets). **E.** RT-qPCR analysis of indicated transcripts following transfection of HEK-293 cells with indicated siRNAs and treatment with 1 μM thapsigargin for 6hr. Relative fold changes are in reference to vehicle- and non-targeting (NT) siRNA-treated cells. Error bars indicate SD (n = 3). Dashed lines indicate log_**2**_(fold change) of ±0.5. PTC+ indicates the use of primers specific to transcript isoforms with validated poison exons (Lareau et al., 2007; Ni et al., 2007). See also Table S2 for statistical comparisons and associated *P* values.

In response to the accumulation of unfolded proteins in the ER, cells activate the integrated stress response (ISR), which restores homeostasis by repressing translation and inducing expression of a battery of stress response genes (Costa-Mattioli & Walter, 2020). Activation of the ISR induces hyperphosphorylation of eIF2ɑ, driving global downregulation of protein synthesis due to impaired eIF2-GTP-Met-tRNAi ternary complex recycling and reduced delivery of the initiator Met-tRNAi to translating ribosomes (Baird & Wek, 2012; Wek, 2018; Young & Wek, 2016). An established effect of ISR-mediated translational repression is corresponding stabilization of well-characterized NMD targets, including several mRNAs encoding factors integral to the activation and resolution of the stress response (Goetz & Wilkinson, 2017). Because of our finding that UPF1_LL_ regulates mRNAs enriched for membrane and ER-associated gene products, we asked how UPF1_LL_ activity responds to ER stress and induction of the ISR (Fig 5B).

To assess the effects of ER stress on UPF1_LL_-dependent regulation, we performed RNA-seq of HEK-293 cells treated with the ER stress-inducing agent thapsigargin. Western blot analysis showed a 2.5-fold increase in eIF2ɑ phosphorylation with thapsigargin treatment (Fig S5B), supporting a robust induction of the ISR. Consistent with previous results (Nickless *et al*, 2014; Li *et al*, 2017), mRNAs up-regulated upon total UPF1 knockdown in HEK-293 cells were on average also up-regulated following 6 hours in 1 μM thapsigargin (Fig 5C), and the magnitude of the increase correlated with the effects of UPF1 total knockdown (interquartile range of log_2_FC of genes >2-fold up-regulated in siUPF1_total_ = −0.02 to 0.56 *vs.* interquartile range of log_2_FC of genes >1.4-fold up-regulated in siUPF1_total_ = −0.07 to 0.36). In sharp contrast, genes up-regulated by UPF1_LL_-specific knockdown exhibited a distinct behavior upon thapsigargin treatment. Rather than increasing, UPF1_LL_ substrates showed on average a reduction in mRNA levels, and this tendency did not vary according to the magnitude of the effect of UPF1_LL_ knockdown (interquartile range of log_2_FC of genes >2-fold up-regulated in siUPF1_LL_ = −0.29 to 0.17 *vs.* interquartile range of log_2_FC of genes >1.4-fold up-regulated in siUPF1_LL_ = −0.26 to 0.19). These results indicate that UPF1_LL_ functions distinctly from that of well-characterized NMD and sustains activity during ER stress and activation of the ISR.

### UPF1_LL_ conditionally remodels NMD target selection during ER stress and ISR induction

To more comprehensively evaluate the role of UPF1_LL_ in promoting the downregulation of select genes during ISR induction, we transfected HEK-293 cells with non-targeting (NT) or UPF1_LL_-specific siRNAs and then treated cells with 1 μM thapsigargin for 6 hours (Fig S5C). In RNA-seq analyses, we identified 606 genes that significantly decreased in abundance with thapsigargin treatment, of which 135 were rescued upon UPF1_LL_ knockdown (Fig 5D). Inferred mRNA stability changes using REMBRANDTS software supported that the observed differences in mRNA abundance upon thapsigargin treatment and UPF1_LL_ knockdown were due to changes in mRNA decay (Fig S5D and E). The changes in gene expression caused by UPF1_LL_ depletion were not attributable to differential ISR induction, as previously established stress response genes were comparably up-regulated in response to thapsigargin treatment following NT and UPF1_LL_-specific knockdown (Fig S5F) (Ashburner *et al*, 2000; The Gene Ontology Consortium, 2019). Moreover, thapsigargin treatment did not alter the relative levels of UPF1_LL_ and UPF1_SL_ mRNAs (Fig S5G), indicating that UPF1_LL_ activity in ER stress was likely due to activity of the existing population of protein rather than a consequence of altered UPF1 splicing upon thapsigargin treatment.

Our finding that UPF1_LL_ has the potential to bind and regulate transcripts normally insensitive to NMD (Fig 4) led us to ask whether genes down-regulated by UPF1_LL_ during ISR induction included substrates beyond those identified as UPF1_LL_ targets under normal cellular conditions (Fig 1). Of the 135 genes downregulated by UPF1_LL_ upon thapsigargin treatment, 49 genes (36%) were unique to the population of UPF1_LL_ targets down-regulated during ISR induction, while 86 genes were identified as UPF1_LL_ targets under both normal and stress conditions. Thus, our RNA-seq and quantitative RT-PCR experiments indicate that UPF1_LL_ activity is maintained or enhanced when cells are subjected to ER stress conditions that inhibit well-characterized NMD events (Fig 5E). In addition to mRNAs that are constitutively regulated by UPF1_LL_ under both normal and stress conditions, these experiments identify a second population of conditionally targeted mRNAs, defined as mRNAs that are not detectably regulated by UPF1_LL_ under normal cellular conditions but undergo UPF1_LL_-dependent downregulation in ER stress. Taken together, these data support a model in which expression of dual UPF1_SL_ and UPF1_LL_ isoforms enable conditional remodeling of NMD target selection in response to ISR induction.

### UPF1_LL_ activity is enhanced by translational repression

We have identified that endogenous UPF1_LL_ regulates the abundance of a population of transcripts that undergo enhanced downregulation during ER stress and induction of the ISR. Because NMD requires detection of in-frame stop codons, NMD target susceptibility is strongly sensitive to changes in the location and frequency of translation initiation and termination. In addition to modulation of initiation via eIF2ɑ phosphorylation in ER stress (Goetz & Wilkinson, 2017), perturbation of translation elongation by selective inhibitors (e.g. cycloheximide and puromycin) inhibits the decay of well-characterized NMD targets (Carter *et al*, 1995). We therefore asked whether translational repression would promote UPF1_LL_ activity outside of the context of the ISR (Fig 6A).

**Figure 6.**
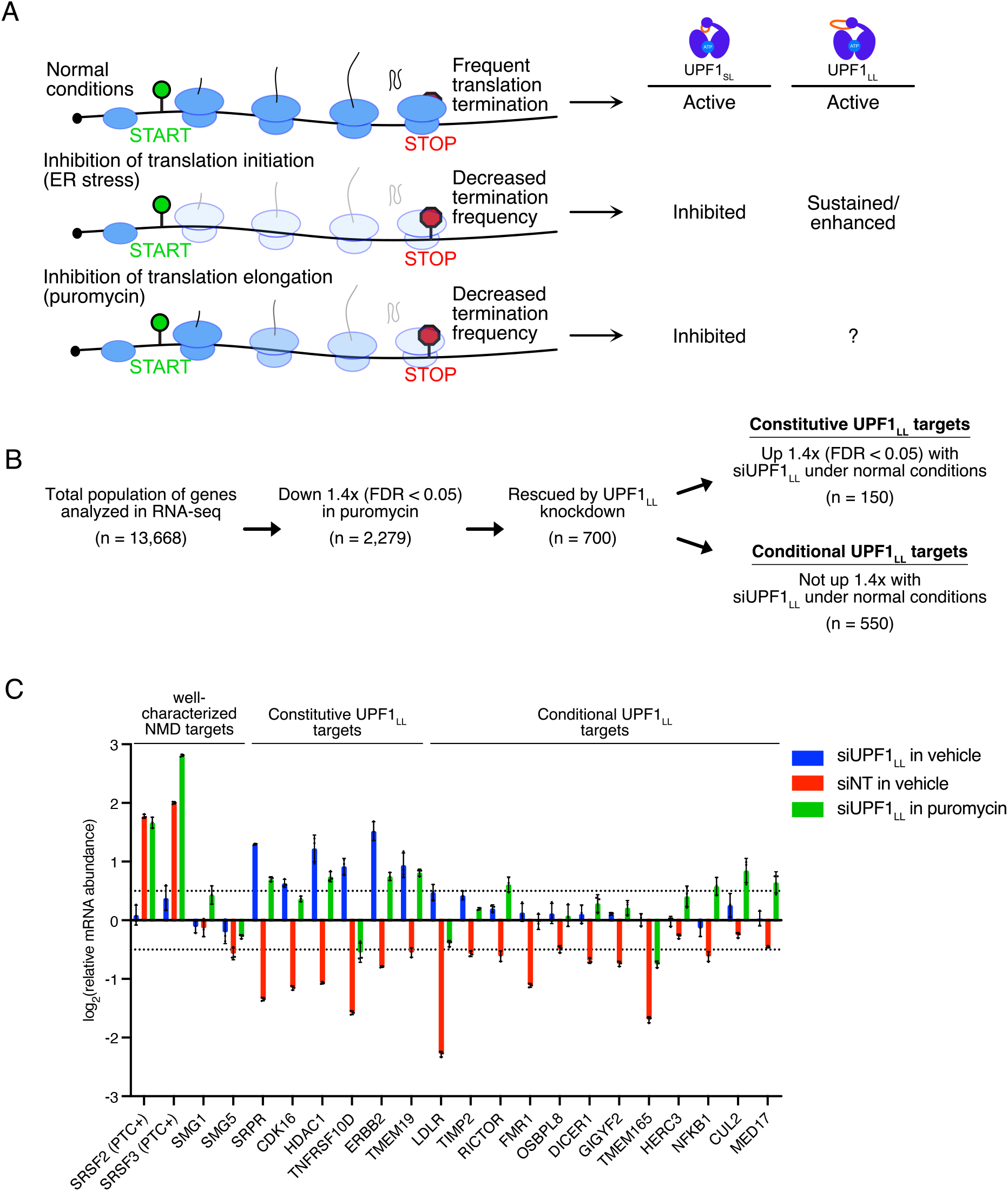
UPF1_LL_ activity is enhanced by translational repression. **A.** Scheme for predicted effect of changes to translation efficiency on UPF1_LL_ activity. Ribosome shading indicates effects of translation initiation and elongation inhibition at different steps in translation. **B.** RNA-seq analysis of HEK-293 cells identifies populations of genes that decreased in abundance with puromycin treatment (50 *μ*g/mL for 4hr) and were rescued by UPF1_LL_-specific knockdown. Indicated are genes that increased in abundance at least 1.4-fold (FDR < 0.05) with UPF1_LL_-specific knockdown under normal conditions (Constitutive UPF1_LL_ targets) and those that did not (Conditional UPF1_LL_ targets). **C.** RT-qPCR analysis of indicated transcripts following transfection of HEK-293 cells with indicated siRNAs and treatment with 50 μg/mL puromycin for 4hr. Relative fold changes are in reference to vehicle- and non-targeting (NT) siRNA-treated cells. Error bars indicate SD (n = 3). Dashed lines indicate logifold change) of ±0.5. PTC+ indicates the use of primers specific to transcript isoforms with validated poison exons (Lareau et al., 2007; Ni et al., 2007). See also Table S2 for statistical comparisons and associated *P* values.

We elected to use the translation elongation inhibitor puromycin because it is widely used to inhibit canonical NMD events and acts through a completely distinct mechanism from the block to initiation caused by eIF2ɑ phosphorylation. Specifically, the ribosome catalyzes the linkage of puromycin to nascent polypeptides, causing chain termination and peptide release (Nathans, 1964). To investigate whether UPF1_LL_ is able to exert sustained or even enhanced post-transcriptional control in response to translation inhibition via distinct mechanisms, we transfected HEK-293 cells with non-targeting (NT) or UPF1_LL_-specific siRNAs and then treated cells with puromycin (50 µg/mL for 4 hours; Fig S6A). Remarkably, RNA-seq analyses revealed 2,279 genes that significantly decreased in abundance with puromycin treatment, of which 700 (31%) were rescued upon UPF1_LL_ knockdown (Fig 6B).

The observation that ISR induction promoted UPF1_LL_ to down-regulate a population of new substrates led us to ask whether we could use puromycin to identify new conditional UPF1_LL_ targets. Strikingly, 550 genes (79%) were unique to the population of UPF1_LL_ targets down-regulated only during puromycin treatment, while the remaining 150 genes (21%) significantly overlapped with those up-regulated with UPF1_LL_ depletion under normal cellular conditions (Fig 6B). Quantitative RT-PCR of select transcripts confirmed the transcriptome-wide RNA-seq results (Fig 6C). Similar to conditions of ER stress, puromycin treatment did not alter UPF1 splicing (Fig S6B), indicating that UPF1_LL_ activity during conditions of impaired translation was likely due to the existing population of UPF1_LL_ protein. These data support that translational repression promotes UPF1_LL_ activity outside of the context of the ISR to conditionally remodel NMD target selection.

Because of the well-established requirement for translation in NMD, we hypothesized that the UPF1 isoform-dependent effects of thapsigargin and moderate puromycin treatment were due to changes in the location and/or frequency of translation termination events (Fig 6A). To test this hypothesis, we treated cells with a titration of puromycin from 25 μg/mL to 400 μg/mL. If UPF1_LL_ activity depends on the infrequent residual translation termination events that occur under puromycin treatment, its activity should be enhanced at low concentrations of puromycin that permit some termination events to persist but be inhibited by high concentrations of puromycin that more efficiently block translation. In line with these expectations, we observed a dose-dependent response, in which downregulation of representative UPF1_LL_ target transcripts FMR1, eIF5A2, ERBB2, and CDK16 was most efficient at lower puromycin concentrations (Fig S6C). Treatment with high concentrations of puromycin did not have a significant effect on the levels of these UPF1_LL_ target mRNAs, consistent with a requirement for translation termination events. Corroborating these findings, we observed globally more efficient downregulation of puromycin-sensitive UPF1_LL_ targets with 25 μg/mL puromycin than 100 μg/mL puromycin in RNA-seq (Fig S6D). Based on these results, we conclude that translation termination is likely required for all NMD events but that changes in translation efficiency can drive downregulation of a novel class of substrates by the UPF1_LL_ isoform.

### Translational repression promotes UPF1_LL_-dependent decay of select mRNAs

Inferred mRNA stability changes using REMBRANDTS software indicated that the observed differences in mRNA abundance upon puromycin treatment and UPF1_LL_ knockdown from the RNA-seq studies were due to corresponding changes in mRNA stability (Fig S6E and F). To directly evaluate the effect of translational repression on promoting the decay of mRNAs by UPF1_LL_, we leveraged the recently established method of Roadblock-qPCR to assess endogenous mRNA stability (Watson *et al*, 2020). In this method, 4-thiouridine (4-SU) is used to label transcripts produced during a 4 hour timecourse. Isolated RNA is treated with N-ethylmaleimide, which covalently labels 4-SU residues, forming a bulky adduct that blocks reverse transcription. RT-qPCR of the remaining unlabeled pool thus allows straightforward quantification of mRNA turnover. Here, HEK-293 cells were transfected with non-targeting (NT) or UPF1_LL_-specific siRNAs and labeled with 4-SU in the absence and presence of puromycin. In this analysis, puromycin treatment stabilized the canonical NMD target of ATF4 (Fig 7A), consistent with previous findings that translational repression inhibits decay of well-characterized NMD targets. In sharp contrast, representative UPF1_LL_ targets CDK16 and TNFRSF10D exhibited significantly shorter half-lives with puromycin treatment, an effect that was dependent upon UPF1_LL_ expression. Together, these data support the conclusion that translational repression promotes the decay of mRNAs by UPF1_LL_.

**Figure 7.**
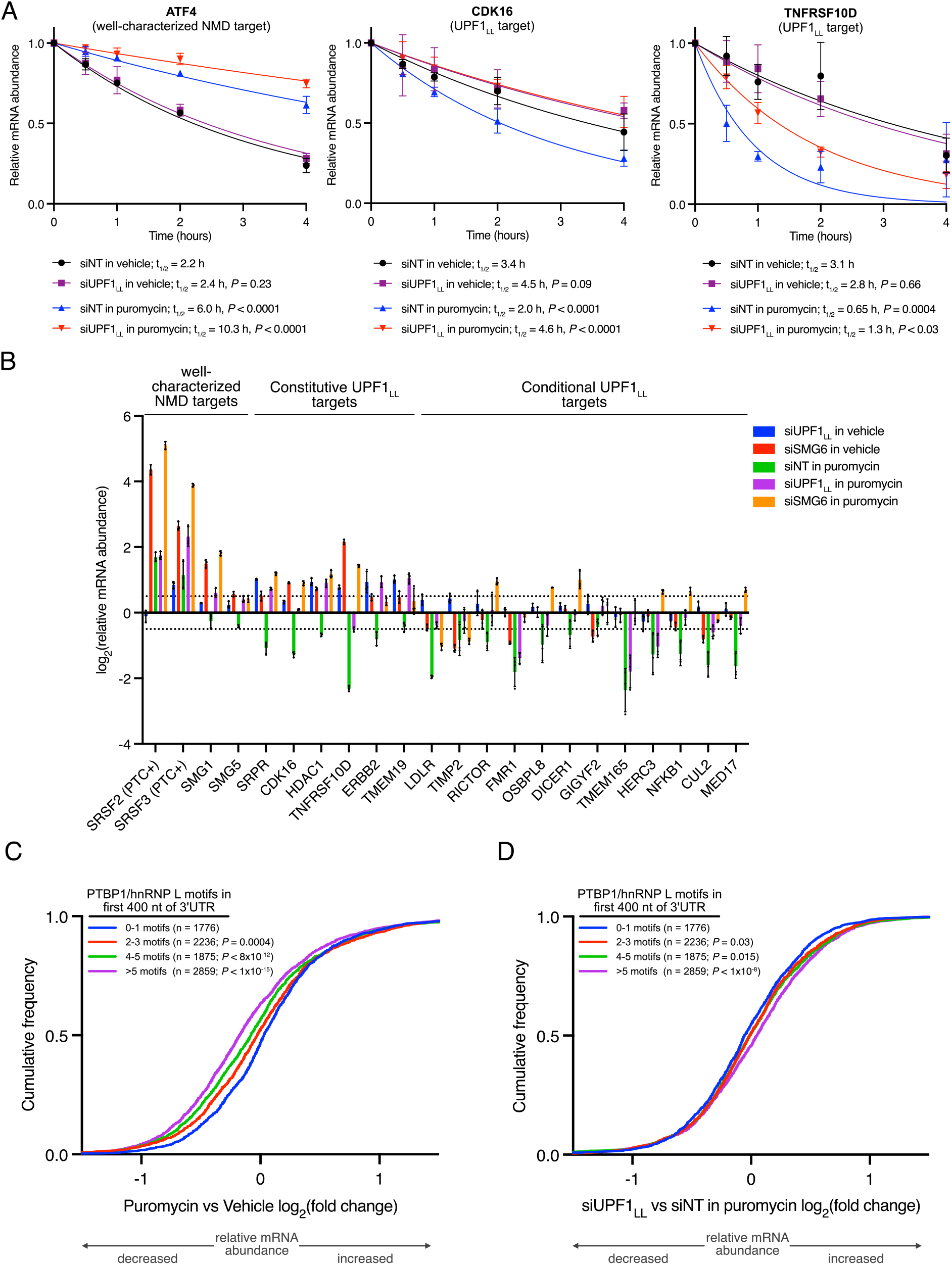
Translational repression promotes the decay of mRNAs by UPF1_LL_ in the NMD pathway and down-regulates normally protected transcripts. **A.** mRNA decay measurements using Roadblock-qPCR (Watson et al., 2020). RNA was isolated from HEK-293 cells at indicated timepoints following transfection with non-targeting (NT) or UPF1 LL-specific siRNAs and treatment with 400 μM 4sU and 50 *μ*g/mL puromycin. Isolated RNA from 4sU-exposed cells was treated with NEM before reverse transcription with oligo-dT primers. Relative levels of the indicated mRNAs at each timepoint were determined by qPCR. mRNA half-lives were estimated by fitting the data to a single-phase exponential decay model. Significance of puromycin treatment was compared to the vehicle control, and significance of UPF1_LL_ knockdown was compared to the NT siRNA in the absence and presence of puromycin treatment. Error bars indicate SD (n = 4). See also Table S2 for 95% confidence intervals associated with mRNA half-lives. **B.** RT-qPCR analysis of indicated transcripts following transfection of HEK-293 cells with indicated siRNAs and treatment with 50 *μ*g/mL puromycin for 4hr. Relative fold changes are in reference to vehicle-treated, non-targeting (NT) siRNA. Error bars indicate SD (n = 3). Dashed lines indicate log_2_(fold change) of ±0.5. PTC+ indicates the use of primers specific to transcript isoforms with validated poison exons (Lareau et al., 2007; Ni et al., 2007). See also Table S2 for statistical comparisons and associated *P* values. **C.** CDF plot of relative mRNA abundance as determined from RNA-seq following treatment of HEK-293 cells with 50 *μ*g/mL of puromycin for 4hr. mRNAs were subdivided by PTBP1 and/or hnRNP L motif density within the first 400 nt of 3’UTR. Statistical significance was determined by K-W test, with Dunn’s correction for multiple comparisons. **D.** CDF plot as in (C), following UPF1_LL_-specific knockdown.

We next asked whether UPF1_LL_-dependent mRNA downregulation during translational repression involved the specialized NMD endonuclease SMG6. To explore this possibility, HEK-293 cells were transfected with non-targeting (NT) or SMG6-specific siRNAs and then treated with puromycin. Quantitative RT-PCR of select transcripts revealed that knockdown of SMG6 significantly rescued the downregulation of UPF1_LL_ targets in puromycin treatment (Fig 7B). This effect was comparable to that observed with UPF1_LL_-specific depletion, supporting the conclusion that select mRNA downregulation during translational repression is due to UPF1_LL_ activity in the NMD pathway. Furthermore, knockdown of SMG6 under normal conditions increased the abundance of well-characterized NMD targets and constitutively-regulated UPF1_LL_ substrates but did not significantly affect the levels of conditionally-regulated UPF1_LL_ substrates. These results provide further evidence that UPF1_LL_ conditionally remodels NMD target selection during translational repression to promote the decay of a new class of substrates.

### NMD-protected mRNAs are down-regulated by UPF1_LL_ during translational repression

Finally, we asked whether the expanded functions of UPF1_LL_ in translational repression are related to its biochemical capability to direct degradation of mRNAs normally protected by PTBP1 and/or hnRNP L. Specifically, we analyzed whether RNAs with stop codon-proximal PTBP1 and hnRNP L motifs are among those susceptible to UPF1_LL_-mediated downregulation upon puromycin treatment. Subdivision of the transcriptome according to PTBP1 and/or hnRNP L motif binding density within the first 400 nt of the 3’UTR revealed that mRNAs with high densities of binding sites for the protective proteins were significantly down-regulated with puromycin treatment relative to mRNAs with low densities of binding sites (Fig 7C). Knockdown of UPF1_LL_ rescued this decrease in mRNA abundance, supporting the conclusion that the downregulation of protected mRNAs during translational repression was dependent upon UPF1_LL_ expression (Fig 7D). Based on these data, we conclude that enhanced UPF1_LL_ activities upon translational repression result in deprotection of normally NMD-insensitive mRNAs.

## Discussion

Here we employ specific depletion, overexpression, and biochemical methods to identify that the mammalian UPF1_LL_ isoform performs distinct functions from that of the major UPF1_SL_ isoform. By depleting only the UPF1_LL_-encoding mRNA, we show that UPF1_LL_ is required for a subset of UPF1-mediated regulation, preferentially targeting mRNAs that encode transmembrane and secreted proteins translated at the ER (Fig 1 and 5). Our transcriptome-wide studies of UPF1_LL_ and UPF1_SL_ RNA binding reveal that UPF1_LL_ has a greater propensity to bind mRNAs normally protected from decay by PTBP1 and hnRNP L (Fig 2). The cellular interaction specificity of UPF1_LL_ is corroborated by its ability to overcome inhibition by PTBP1 *in vitro* (Fig 3), consistent with our previous observation that PTBP1 promotes ATPase-dependent UPF1 dissociation by exploiting the UPF1 regulatory loop (Fritz *et al*, 2020). Overexpression of UPF1_LL_ leads to preferential downregulation of mRNAs with long 3’UTRs that are normally protected by PTBP1 and hnRNP L (Fig 4). These data in sum suggest that UPF1_LL_ has the biochemical capacity to regulate the protected population of mRNAs but that its activities are likely constrained by its relatively low expression level in HEK-293 and many other cell types.

Based on the observation that hundreds of genes regulated by UPF1_LL_ under normal conditions encode proteins trafficked through the ER, we investigated the effects of ER stress on UPF1_LL_ function. In striking contrast to the well-characterized inhibition of NMD by ER stress, we find that UPF1_LL_-dependent regulation is intact or even enhanced (Fig 5). Mechanistically, we find that preferential UPF1_LL_ activity in ER stress can be explained by its ability to function under conditions of translational repression (Fig 6). Moderate inhibition of translation with puromycin causes thousands of genes to be down-regulated, of which approximately one-third are rescued by UPF1_LL_ knockdown. Importantly, these experiments show that UPF1_LL_ is not only required for downregulation of mRNAs identified as substrates under normal conditions (“constitutive” UPF1_LL_ targets), but also regulates additional mRNAs under ER stress and translational repression (“conditional” UPF1_LL_ targets). Providing a mechanism for conditional targeting of normally protected mRNAs, mRNAs that are down-regulated by UPF1_LL_ upon puromycin treatment are enriched for TC-proximal protective protein binding sites (Fig 7). Combined with the inhibition of UPF1_SL_-dependent decay, enhanced UPF1_LL_ activity upon translational repression results in a dramatic and unanticipated shift in NMD target specificity.

The differential RNA-binding properties of UPF1_SL_ and UPF1_LL_ in the presence of ATP and the ability of the protective proteins to drive ATPase-dependent UPF1 dissociation provide a molecular mechanism by which NMD specificity is tuned in response to cellular conditions (Gowravaram *et al*, 2018; Fritz *et al*, 2020). We present a model in which the increased residence time of UPF1_LL_ on RNAs enables it to respond to infrequent translation termination events, while the faster dissociation of UPF1_SL_ from potential NMD targets renders it unable to complete decay when translation is repressed (Fig 8). The ability of UPF1_LL_ to maintain functionality in response to infrequent termination events thus enables a shift in NMD specificity when translation is partially inhibited. As translation efficiency decreases, the relative activity of UPF1_LL_ exceeds that of UPF1_SL_, allowing downregulation of new substrate mRNAs, including those normally protected from decay by PTBP1 and hnRNP L. The rate-limiting step(s) downstream of UPF1 binding to potential NMD targets remain to be clearly defined, but an attractive possibility is that persistent binding of UPF1_LL_ to mRNAs promotes its phosphorylation by SMG1, readying UPF1_LL_ to scaffold assembly of productive decay complexes (Durand *et al*, 2016).

**Figure 8.**
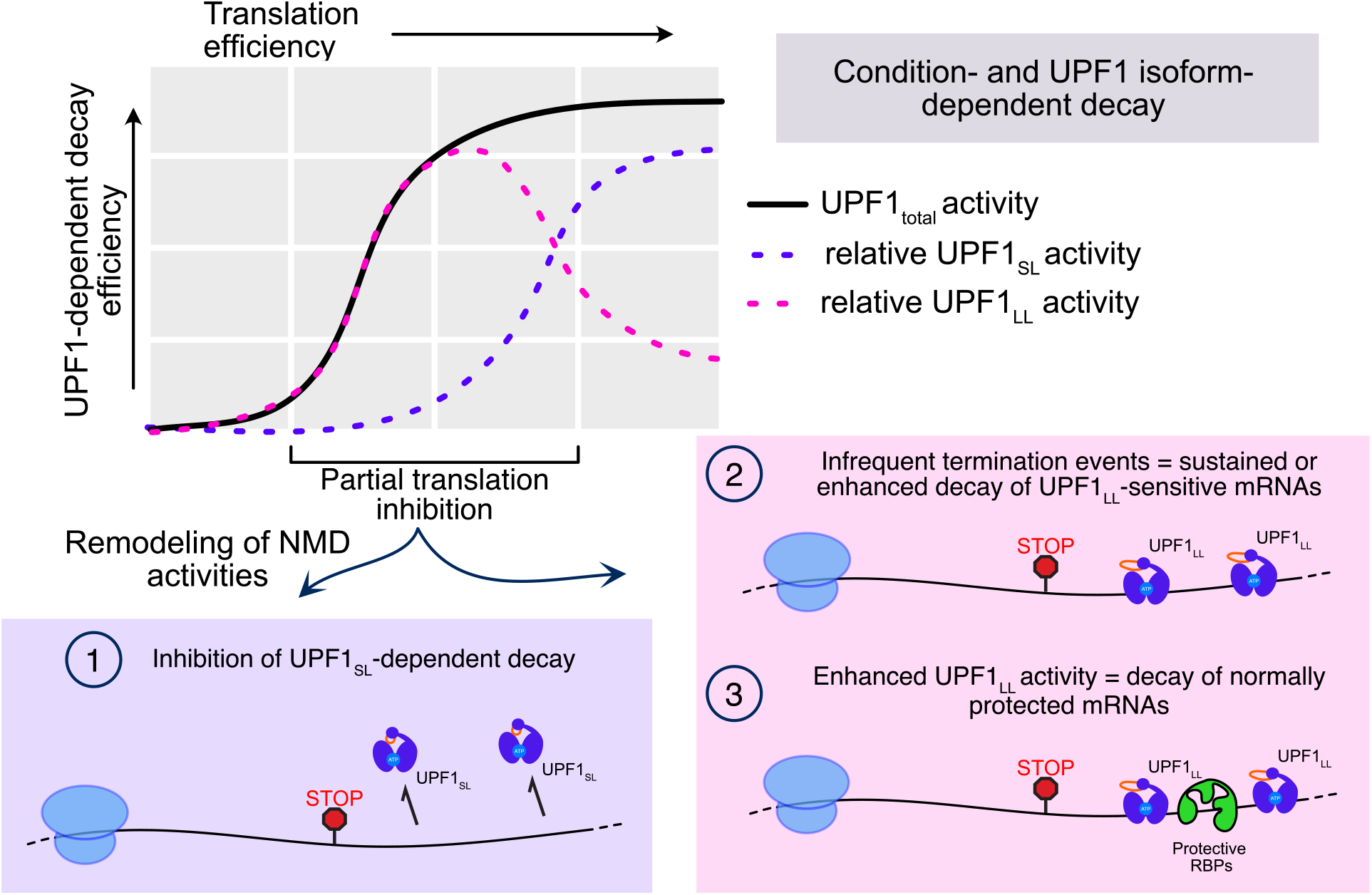
Translation efficiency and UPF1 isoform expression conditionally alter target susceptibility by the NMD pathway. UPF1_LL_ retains activity under conditions of partial translation repression that efficiently inhibit UPF1_SL_· As translation efficiency decreases, cellular NMD specificity shifts, causing enhanced targeting of constitutive UPF1_LL_-regulated mRNAs and conditional UPF1_LL_-dependent substrates. mRNAs conditionally sensitive to UPF1_LL_ include those normally protected by PTBP1 or hnRNP L.

The idea that the mammalian NMD pathway consists of multiple branches with distinct factor requirements and substrate specificities has been proposed by several groups, but the underlying mechanisms and regulatory roles of NMD branching are poorly understood (Huang *et al*, 2011; Chan *et al*, 2007; Yi *et al*, 2020). Our identification of differential activities of UPF1_SL_ and UPF1_LL_ is an unforeseen example of NMD pathway branching, which can be controlled at the cellular level by changes in the abundance of protective RBPs, translation, or UPF1 splicing. Because altered translation efficiency and induction of cellular stress pathways are pervasive in cancer, genetic disease, and infection, the expanded scope of UPF1_LL_-dependent decay has far-reaching implications for the physiological roles of the mammalian NMD pathway in health and disease.

In particular, the ability of UPF1_LL_ to target specific mRNAs in response to translational repression positions NMD to function as a mechanism to reset the transcriptome upon cellular stress. Translational repression via eIF2α phosphorylation is a central feature of responses to diverse cellular insults, including nutrient deprivation, oncogene activation, viral infection, hypoxia, and accumulation of unfolded proteins (Pakos-Zebrucka *et al*, 2016). We show that induction of ER stress with thapsigargin induces UPF1_LL_-dependent downregulation of hundreds of transcripts, a population particularly enriched for mRNAs encoding proteins trafficked through the ER. The extent to which UPF1_LL_ complements or collaborates with IRE1-mediated mRNA decay or the recently identified ER-associated NMD factor NBAS will require further study (Hollien & Weissman, 2006; Longman *et al*, 2020), but our data suggest that UPF1_LL_ may help to relieve proteotoxic stress by reducing the abundance of transcripts encoding proteins with complex biosynthetic needs.

Notably, we find that UPF1_LL_ is able to conditionally regulate several proteins of central importance in cancer and other diseases, including fragile X mental retardation 1 (FMR1), the low-density lipoprotein receptor (LDLR), and oncogenes PTEN, EIF5A2, and ERBB2, among others (Dockendorff & Labrador, 2019; Goldstein & Brown, 2009; Song *et al*, 2012; Mathews & Hershey, 2015; Harbeck *et al*, 2019). Along with EIF5A2 and FMR1, several additional UPF1_LL_ targets that undergo enhanced downregulation upon translational repression are themselves important regulators of translation and mRNA decay, including the signal recognition particle receptor SRPR, the major mRNA decapping enzyme DCP2, and the miRNA-processing endonuclease DICER1 (Michlewski & Cáceres, 2019; Akopian *et al*, 2013; Mugridge *et al*, 2018). Even in the absence of stress, most cells and tissues *in vivo* likely have lower basal rates of translation than those attained in exponentially growing transformed cell lines. Based on our findings, understanding how NMD shapes gene expression in diverse tissue types in health and disease will require not just analysis of NMD targets characterized in transformed cells, but also transcripts that are conditionally targeted in response to changing translational states.

## Materials and methods

### UPF1 isoform nomenclature

UPF1_SL_ refers to the protein encoded by Ensembl transcript ID ENST00000262803.9 and UPF1_LL_ by Ensembl transcript ID ENST00000599848.5. These isoforms are also referred to as UPF1 isoform 2 and UPF1 isoform 1, respectively (Gowravaram *et al*, 2018; Nicholson *et al*, 2014).

### *In vitro* helicase assays

Unwinding assays were performed as previously described in detail (Fritz *et al*, 2020). Briefly 75 nM of the pre-assembled RNA duplex substrate (described below) was combined with 1x unwinding reaction buffer (10 mM MES pH 6.0, 50 mM KOAc, 0.1 mM EDTA), 2 mM MgOAc, and 1 unit RNasin in a well of a half-area, black, flat-bottom 96-well plate (Corning 3993). PTBP1 (80 nM) was then added, the sample mixed, and incubated at room temperature for 10 minutes. Then, UPF1 (80 nM) was added, the sample mixed, and incubated at room temperature for 10 minutes. Finally, BHQ1 quencher (0.56 μM final; GTGTGCGTACAACTAGCT / 3BHQ_1) and 2 mM ATP was added to initiate the unwinding reaction. An Infinite^R^ F200 Pro microplate reader and associated i-control™ 1.9 software (Tecan) was used to monitor Alexa Fluor 488 fluorescence every 10 seconds for 20 minutes at 37°C. Measured fluorescence intensities were normalized to the zero timepoint for each condition to obtain relative fluorescence values. Four technical replicates for each condition were obtained at the same time. The decrease in fluorescence caused by UPF1 unwinding in the absence of PTBP1 at the end of each 20 minute timecourse was used to calculate the time to 50% maximal undwinding and relative total unwinding for each condition. Because measurements were taken at 10s intervals, the earliest time at which 50% maximal unwinding was observed is indicated on each graph.

To generate the RNA duplex, a DNA oligo template (5’ TAATACGACTCACTATAGGGACACAAAACAAAAGACAAAAACACAAAACAAAAGACAAAAACA CAAAACAAAAGACAAAAAGCCTCTCCTTCTCTCTGCTTCTCTCTCGCTGTGTGCGTACAACTAGCT 3’) was PCR-amplified and then *in vitro* transcribed using the MEGAshortscript™ T7 Transcription kit (Invitrogen). An 11:7 ratio of the helicase RNA substrate to 5’ Alex Fluor 488 fluorescent oligo strand (Alexa Fluor 488/ AGCTAGTTGTACGCACAC) was incubated with 2 mM MgOAC and 1x unwinding reaction buffer at 95°C for 3 minutes and 30 seconds and then slowly cooled to 30°C. All DNA oligos were obtained from Integrated DNA Technologies. The 5’ fluorescent oligo probe and 3’ BHQ1 quencher were acquired as RNase-free, HPLC-purified.

Cloning, expression, and purification of recombinant UPF1_LL_ΔCH, UPF1_SL_ΔCH, and PTBP1 were conducted as previously described (Fritz *et al*, 2020; Gowravaram *et al*, 2018). In this study, UPF1_LL_ΔCH was purified in the same manner as UPF1_SL_ΔCH (Fritz *et al*, 2020).

### Mammalian cell lines and generation of CLIP-UPF1 expression lines

HEK-293 cells used in the endogenous UPF1_LL_ knockdown and RNA-seq experiments were received from ATCC (CRL-3216) and maintained at 37°C and 5% CO_2_ in DMEM with 10% FBS (Gibco) and 1% pen/strep. Stable integration of CLIP-UPF1_SL_ into human Flp-InTM T-RExTM-293 cells (Invitrogen) and subsequent maintenance of this stable line was previously described (Kishor *et al*, 2020). CLIP-UPF1_LL_ expression lines were generated and maintained in an identical manner. Total UPF1 and SMG6 depletion studies followed by RT-qPCR as well as the Roadblock-qPCR experiments were performed using HEK-293 cells maintained as described above. Total UPF1 depletion and RNA-seq was conducted using parental Flp-In™ T-REx™-293 cells (Invitrogen) maintained according to manufacturer instructions.

### Endogenous UPF1 or SMG6 depletion by siRNA

The following siRNAs were used to deplete indicated UPF1 isoforms or the SMG6 endonuclease: total UPF1 (Forward sequence: 5’ CUACCAGUACCAGAACAUA 3’; Reverse sequence: 5’ UAUGUUCUGGUACUGGUAG 3’); UPF1_LL_ (Forward sequence: 5’ GGUAAUGAGGAUUUAGUCA 3’; Reverse sequence: 5’ UGACUAAAUCCUCAUUACC 3’); SMG6 (Forward sequence: 5’ GCUGCAGGUUACUUACAAG 3’; Reverse sequence: 5’ CUUGUAAGUAACCUGCAGC 3’; (Durand *et al*, 2016)).

### CLIP-UPF1 overexpression RIP-seq and RNA-seq

CLIP-UPF1_SL_ overexpression RIP-seq and RNA-seq sample preparation were previously reported (Kishor *et al*, 2020); CLIP-UPF1_LL_ datasets were generated in parallel. Briefly, CLIP-UPF1 stable cell lines or a GFP-expressing control line were seeded in 6 × 15 cm plates and then treated with 200 ng/mL doxycycline hyclate (Sigma) for 48 hours to induce CLIP-UPF1 expression. Cells were harvested 48 hours post-induction at 80-85% confluency and whole cell lysate generated by the freeze/thaw method as previously described (Hogg & Collins, 2007a, 2007b; Hogg & Goff, 2010; Fritz *et al*, 2018). Equilibrated cell extracts, reserving 1/10th for downstream input analysis, were then combined with 10 μM CLIP-Biotin (New England Biolabs) and rotated (end-over-end) for 1 hour at 4°C to allow CLIP-UPF1 to react with the CLIP-Biotin substrate and yield covalently biotinylated protein. Unbound CLIP-Biotin was subsequently removed by passing the samples through Zeba™ Spin Desalting Columns, 40K MWCO, 2 mL (Thermo Scientific) according to manufacturer instructions. Buffer exchange was performed with HLB-150 supplemented with 1 mM DTT and 0.1% NP-40. Pre-washed Dynabeads™ MyOne™ Streptavidin T1 (Invitrogen) were then added and the samples rotated (end-over-end) for 1 hour at 4°C to immobilize the biotin-bound CLIP-UPF1 complexes. The samples were then washed three times with 500 μL of HLB-150 supplemented with 1 mM DTT and 0.1% NP-40 and one one-hundredth reserved for downstream Western blot analysis of CLIP-UPF1 pull-down efficiency. The remainder was combined with TRIzol™ (Invitrogen) and RNA was isolated according to manufacturer instructions. DNase-treatment was subsequently performed using RQ1 RNase-Free DNase (Promega), and RNA was isolated by acid phenol chloroform extraction according to standard protocol. This resulted in approximately 1 μg of total RNA that was then subjected to high-throughput sequencing. In parallel, the reserved input lysate (approximately 3 μg of total RNA) was also sent for high-throughput sequencing. A total of three biological replicates from each condition were processed. Sequencing libraries were prepared from input and bound RNA using the Illumina TruSeq Stranded Total RNA Human kit and sequenced on an Illumina HiSeq 3000 instrument.

To validate select transcripts by RT-qPCR, an equivalent volume of reserved RNA (3 μL) was used as a template for cDNA synthesis using the Maxima First Strand cDNA synthesis kit for RT-qPCR (Thermo Scientific) according to manufacturer instructions. The resulting cDNA was diluted with nuclease-free water and subsequently analyzed by qPCR using iTaq Universal SYBR Green Supermix (BioRad) on a Roche LightCycler 96 instrument (Roche). Sequences for gene-specific primers used for amplification are listed in Table S5. For input samples, relative fold change was determined by calculating 2^−ΔΔCT^ values using GAPDH for normalization. For pull-down samples, relative fold enrichment was determined by dividing the C_q_ value of the pull-down by its corresponding input, multiplying by 100 and then normalizing to the relative recovery of the SMG1 transcript.

For assessment of CLIP-UPF1 expression and pull-down efficiency by Western blot, reserved input and IP samples were run on a NuPage™ 4-12% Bis-Tris Protein Gel (Invitrogen) using MOPS buffer according to manufacturer instructions and subsequently transferred to a nitrocellulose membrane according to the NuPage™ manufacturer’s protocol (Invitrogen). Membranes were incubated with a blocking buffer for fluorescent Western blotting (Rockland) for 1 hour at room temperature and then incubated overnight at 4°C with the indicated primary antibody. Primary antibodies used were: anti-RENT1 (goat polyclonal, Bethyl, A300-038A, 1:1000) and anti-β-actin (mouse monoclonal, Cell Signaling, #3700, 1:1000). Membranes were subsequently washed three times with 1xTBS supplemented with 0.1% Tween-20 and then incubated with the appropriate secondary antibody for 1 hour at room temperature. Secondary antibodies used were: anti-goat IgG (H&L) Antibody DyLight™ 680 Conjugated (Rockland, 605-744-002, 1:10,000) and anti-mouse IgG (H&L) Antibody DyLight™ 680 Conjugated (Rockland, 610-744-124, 1:10,000). Membranes were washed three times with 1xTBS supplemented with 0.1% Tween-20 and then two times with 1xTBS. Western blot images were obtained on an Amersham Typhoon imaging system (GE Healthcare Life Sciences) and quantified using ImageStudio software (LI-COR Biosciences).

For total UPF1 depletion and rescue with CLIP-UPF1, 3×10^5^ cells from the CLIP-UPF1 stable cell lines, which were engineered as resistant to the described total UPF1 siRNA, or the GFP-expressing control line were reverse transfected with 40 nM of a non-targeting or UPF1-specific siRNA (described above) using Lipofectamine RNAiMAX according to manufacturer instructions. The next day, cells were treated with 200 ng/mL doxycycline hyclate (Sigma) for 48 hours to induce expression of CLIP-UPF1. Cells were harvested 48 hours post-induction and total RNA was isolated using TRIzol™ (Invitrogen) according to manufacturer instructions. DNase-treatment was subsequently performed using RQ1 RNase-Free DNase (Promega) and RNA was isolated by acid phenol chloroform extraction according to standard protocol. RT-qPCR was performed as described above.

### Total UPF1 depletion and RNA-seq

3×10^5^ Flp-In™ T-REx™-293 cells (Invitrogen) were reverse transfected with 40 nM of a non-targeting or UPF1-specific siRNA that targets both UPF1 isoforms (described above) using Lipofectamine RNAiMAX according to manufacturer instructions. Seventy-two hours post-siRNA transfection, total RNA was isolated using TRIzol™ (Invitrogen) according to manufacturer instructions. DNase-treatment was subsequently performed using RQ1 RNase-Free DNase (Promega) and RNA was isolated by acid phenol chloroform extraction according to standard protocol. A total of 2 μg RNA was then subjected to high-throughput sequencing. Three replicates were processed for each condition. Sequencing libraries were prepared using the Illumina TruSeq Stranded Total RNA Human kit and sequenced on an Illumina NovaSeq 6000 instrument.

### Total UPF1 and SMG6 depletion with RT-qPCR

3×10^5^ HEK-293 cells were reverse transfected with 40 nM of a non-targeting, total UPF1 or SMG6-specific siRNA (described above) using Lipofectamine RNAiMAX according to manufacturer instructions. Forty-eight hours post-siRNA transfection, cells were replated at a density of 5×10^5^ cells per well of a 6-well plate. This step was critical to achieve 70-75% confluency on the day of drug treatment (if applicable) and cell harvest. The next day, cells were directly harvested or treated with 50 μg/mL puromycin (Sigma) for 4 hours. A total of three replicates were generated for each condition. Total RNA was then isolated using TRIzol™ (Invitrogen) according to manufacturer instructions. DNase-treatment was subsequently performed using RQ1 RNase-Free DNase (Promega), and RNA was isolated by acid phenol chloroform extraction according to standard protocol. For RT-qPCR, 500 ng of total RNA was used as input for cDNA synthesis using the Maxima First Strand cDNA synthesis kit for RT-qPCR (Thermo Scientific) according to manufacturer instructions. The resulting cDNA was diluted with nuclease-free water and subsequently analyzed by qPCR using iTaq Universal SYBR Green Supermix (BioRad) on a Roche LightCycler 96 instrument (Roche). Sequences for gene-specific primers used for amplification are listed in Table S5. Relative fold changes were determined by calculating 2^−ΔΔCT^ values using GAPDH for normalization.

### Endogenous UPF1_LL_ knockdown with puromycin or thapsigargin treatment and RNA-seq

3×10^5^ HEK-293 cells were reverse transfected with 40 nM of a non-targeting or UPF1_LL_-specific siRNA (described above) using Lipofectamine RNAiMAX according to manufacturer instructions. Forty-eight hours post-siRNA transfection, cells were replated at a density of 5×10^5^ cells per well of a 6-well plate. The next day, cells were treated with vehicle control, 25, 50, or 100 μg/mL puromycin (Sigma) for 4 hours, or 1 μM thapsigargin for 6 hours. A total of three replicates were generated for each condition. Total RNA was then isolated using the RNeasy Plus Mini Kit (QIAGEN). Sequencing libraries were prepared from 2 μg total RNA using the Illumina TruSeq Stranded Total RNA Human kit and sequenced on an NovaSeq 6000 instrument. For RT-qPCR validation of changes in target gene expression, 500 ng of total RNA was used as input for cDNA synthesis using the Maxima First Strand cDNA synthesis kit for RT-qPCR (Thermo Scientific) according to manufacturer instructions. The resulting cDNA was diluted with nuclease-free water and subsequently analyzed by qPCR using iTaq Universal SYBR Green Supermix (BioRad) on a Roche LightCycler 96 instrument (Roche). Sequences for gene-specific primers used for amplification are listed in Table S5. Relative fold changes were determined by calculating 2^−ΔΔCT^ values using GAPDH for normalization.

### Roadblock-qPCR to measure endogenous mRNA stability

mRNA decay measurements were determined using Roadblock-qPCR as previously described (Watson *et al*, 2020) but with the following adaptations. 3×10^5^ HEK-293 cells were reverse transfected with 40 nM of a non-targeting or UPF1_LL_-specific siRNA (described above) using Lipofectamine RNAiMAX according to manufacturer instructions. Forty-eight hours post-siRNA transfection, cells were replated at a density of 5×10^5^ cells per well of a 6-well plate. The next day, cells were treated with 400 μM 4-thiouridine (4sU; Cayman Chemical) and vehicle control or 50 μg/mL puromycin (Sigma) for a total of 4 hours. Cells were harvested at indicated timepoints and total RNA was isolated using TRIzol™ (Invitrogen) according to manufacturer instructions, but with the addition of 1 mM (final) DTT to the isopropanol precipitation in order to maintain 4sU in a reduced state (Schofield *et al*, 2018). Isolated RNA from 4sU-exposed cells was treated with 48 mM N-ethylmaleimide (NEM; Sigma) as described by Watson *et al* and then purified using RNAClean XP beads (Beckman Coulter) according to manufacturer instructions. A total of 1 μg RNA was used as input for cDNA synthesis with oligo dT_18_ primers and Protoscript II reverse transcriptase (New England BioLabs) according to manufacturer instructions. The resulting cDNA was diluted with nuclease-free water and subsequently analyzed by qPCR using iTaq Universal SYBR Green Supermix (BioRad) on a Roche LightCycler 96 instrument (Roche). Sequences for gene-specific primers used for amplification are listed in Table S5. Relative fold changes were determined by calculating 2^−ΔΔCT^ values using time 0 as the reference and GAPDH for normalization. mRNA half-lives were estimated by fitting the data to a single-phase exponential decay model using GraphPad Prism 9.1.0.

### Western for phospho-eIF2a

HEK-293 cells treated with 1 μM thapsigargin were lysed in 1X Passive Lysis Buffer (Promega) supplemented with Halt™ Protease and Phosphatase Inhibitor Cocktail (ThermoScientific) according to manufacturer instructions. A total of 5 μg protein was run on a NuPage™ 4-12% Bis-Tris Protein Gel (Invitrogen) using MOPS buffer according to manufacturer instructions and subsequently transferred to a nitrocellulose membrane according to the NuPage™ manufacturer's protocol (Invitrogen). Detection of phospho and total eIF2ɑ was performed as previously described (Young-Baird *et al*, 2020).

### Semiquantitative PCR to detect UPF1 isoform ratios

Generated cDNA (1/40^th^) from RNA-seq samples was used as input for PCR amplification with Phusion High-Fidelity DNA polymerase (New England Biolabs) and UPF1 specific primers that flank the regulatory loop sequence (Forward: 5’ AACAAGCTGGAGGAGCTGTGGA 3’; Reverse: 5’ ACTTCCACACAAAATCCACCTGGAAGTT 3’). The PCR cycling conditions used were: an initial denaturation at 98°C for 30 seconds and then 22 cycles of 98°C for 10 seconds, 63°C for 30 seconds, 72°C for 15 seconds. PCR products were then run on a 8% Novex™ TBE gel (Invitrogen) according to manufacturer instructions and subsequently stained with SYBR^®^ Gold Nucleic Acid Stain (Invitrogen). Images were obtained on an Amersham Typhoon imaging system (GE Healthcare Life Sciences) and quantified using ImageStudio software (LI-COR Biosciences).

### Gene-level differential expression analysis

For analysis of RNA-seq data from HEK-293 cells treated with non-targeting, anti-UPF1_total_, or anti-UPF1_LL_ siRNAs, raw fastq reads from the NovaSeq 6000 platform were trimmed with fastp (Chen *et al*, 2018), with the parameters --detect_adapter_for_pe and --trim_poly_g. Trimmed reads were aligned with HISAT2 to the hg19/GRCh37 genome and transcriptome index provided by the authors (Kim *et al*, 2019). For gene-level differential expression analysis, reads mapping to Ensembl GRCh37 release 75 gene annotations were quantified with featureCounts (Liao *et al*, 2014), and differential gene expression was analyzed using limma/voom, as implemented by the Degust server (Ritchie *et al*, 2015; Powell, 2015).

### Isoform-level differential expression analysis

For isoform-level differential expression analysis, trimmed reads were quantified against a custom HEK-293 transcriptome index, prepared with Stringtie and TACO as described (Kishor *et al*, 2018; Niknafs *et al*, 2017; Pertea *et al*, 2015), using kallisto software with parameters --bias -b 1000 -t 16 --single --rf-stranded -l 200 -s 20 (Bray *et al*, 2016). Differential transcript expression analysis was performed using RUVSeq and edgeR (Risso *et al*, 2014; Robinson *et al*, 2009). Normalization and batch correction was performed with the RUVg function, based on transcripts with invariant expression among all samples. The edgeR TMM method was used to obtain normalized differential expression values and to calculate FDRs.

IsoformSwitchAnalyzeR was used to annotate PTCs in the custom HEK-293 transcriptome, as described (Vitting-Seerup & Sandelin, 2019; Kishor *et al*, 2018). Genes represented by at least one transcript predicted to contain a TC within 50 nt of the final exon junction or in the last exon and at least one transcript predicted to contain a PTC, defined as a TC more than 50 nt upstream of the final exon junction, were selected for analysis of differential isoform usage upon total UPF1 and UPF1_LL_-specific knockdown. Differential isoform usage was calculated for the most abundant PTC and non-PTC isoforms of each gene using IsoformSwitchAnalyzer and the DEXSeq package (Anders *et al*, 2012; Vitting-Seerup & Sandelin, 2019). As a secondary metric capturing NMD targets that do not contain PTCs, NMD-sensitive transcripts were classified on the basis of a log2 fold change of 0.5 or greater in previous RNA-seq studies of total UPF1 knockdown in HEK-293 cells (GSE105436) (Baird *et al*, 2018). Alternative splicing was analyzed using rMATS 4.0.1 (Shen *et al*, 2014), and Sashimi plots were generated using Integrative Genomics Viewer 2.8.2 (Thorvaldsdóttir *et al*, 2013).

### Analysis of 3’UTR length and protective protein motifs

The most abundant transcript isoform from each gene, as determined by quantification with kallisto as above, was used for analysis of 3’UTR length and PTBP1 and hnRNP L binding motif positions and frequencies. PTBP1 and hnRNP L binding motif position-specific scoring matrices were downloaded from the RBPmap database and used for motif finding in 3’UTRs derived from the custom HEK-293 transcriptome with HOMER, as described (Paz *et al*, 2014; Heinz *et al*, 2010; Kishor *et al*, 2018).

### RIP-seq analysis

Raw fastq reads from CLIP-UPF1 overexpression RNA-seq and RIP-seq data were trimmed with Cutadapt using the following parameters: --times 2 -e 0 -O 5 --quality-cutoff 6 -m 18 -a AGATCGGAAGAGCACACGTCTGAACTCCAGTCAC -A AGATCGGAAGAGCGTCGTGTAGGGAAAGAGTGTAGATCTCGGTGGTCGCCGTATCATT -b AAAAAAAAAAAAAAAAAAAAAAAAAAAAAAAAAAAAAAAAAAAAAAAAAA -b TTTTTTTTTTTTTTTTTTTTTTTTTTTTTTTTTTTTTTTTTTTTTTTTTT (Martin, 2011). Trimmed reads were quantified using kallisto software with parameters --bias -b 1000 -t 16 --single --rf-stranded −l 200 -s 20 (Bray *et al*, 2016). RIP-seq enrichment values were obtained by dividing TPM values from IP samples by TPM values from input samples.

### GTEx data analysis

GTEx data were downloaded from the GTEx Portal on 11/11/2020. To determine the relative representation of UPF1_LL_ and UPF1_SL_ mRNA isoforms, transcript TPM values for transcript ENST00000599848.5 (UPF1_LL_) were divided by the total TPM values derived from transcripts ENST00000599848.5 (UPF1_LL_) and ENST00000262803.9 (UPF1_SL_). GTEx samples were assigned to the indicated tissue types using the sample attributes provided in GTEx Analysis v8.

### Data availability

CLIP-UPF1 overexpression RIP-seq and RNA-seq data are available from the NCBI GEO database with accession number GSE134059. The endogenous UPF1_LL_ knockdown RNA-seq data, including puromycin or thapsigargin treatment, are available with accession number GSE162699. Total UPF1 knockdown RNA-seq is available with accession number GSE176197.

## Acknowledgements

We thank Sutapa Chakrabarti for the UPF1_LL_ΔCH recombinant expression plasmid and members of the Hogg lab and Nicholas R. Guydosh for critical reading of the manuscript. We are grateful to Sandy Mattijssen and Richard J. Maraia for helpful discussion and to Sara K. Young-Baird for thapsigargin reagents, helpful discussion, and assistance with phospho-eIF2a immunoblotting. High-throughput sequencing was conducted by Yan Luo and Poching Liu in the NHLBI DNA Sequencing and Genomics Core. The Genotype-Tissue Expression (GTEx) Project was supported by the Common Fund of the Office of the Director of the National Institutes of Health, and by NCI, NHGRI, NHLBI, NIDA, NIMH, and NINDS. This work was supported by the Intramural Research Program, National Institutes of Health, National Heart, Lung, and Blood Institute and utilized the computational resources of the NIH HPC Biowulf cluster (http://hpc.nih.gov).

## Author contributions

S.E.F. and J.R.H. conceived and designed the study, analyzed and interpreted the data, and wrote the manuscript. S.E.F. acquired the UPF1 RIP-seq and RNA-seq data and performed associated cellular studies. S.R. acquired the unwinding data. C.D.W. performed the total UPF1 knockdown. All authors read and approved the manuscript.

---

**Figure S1.**
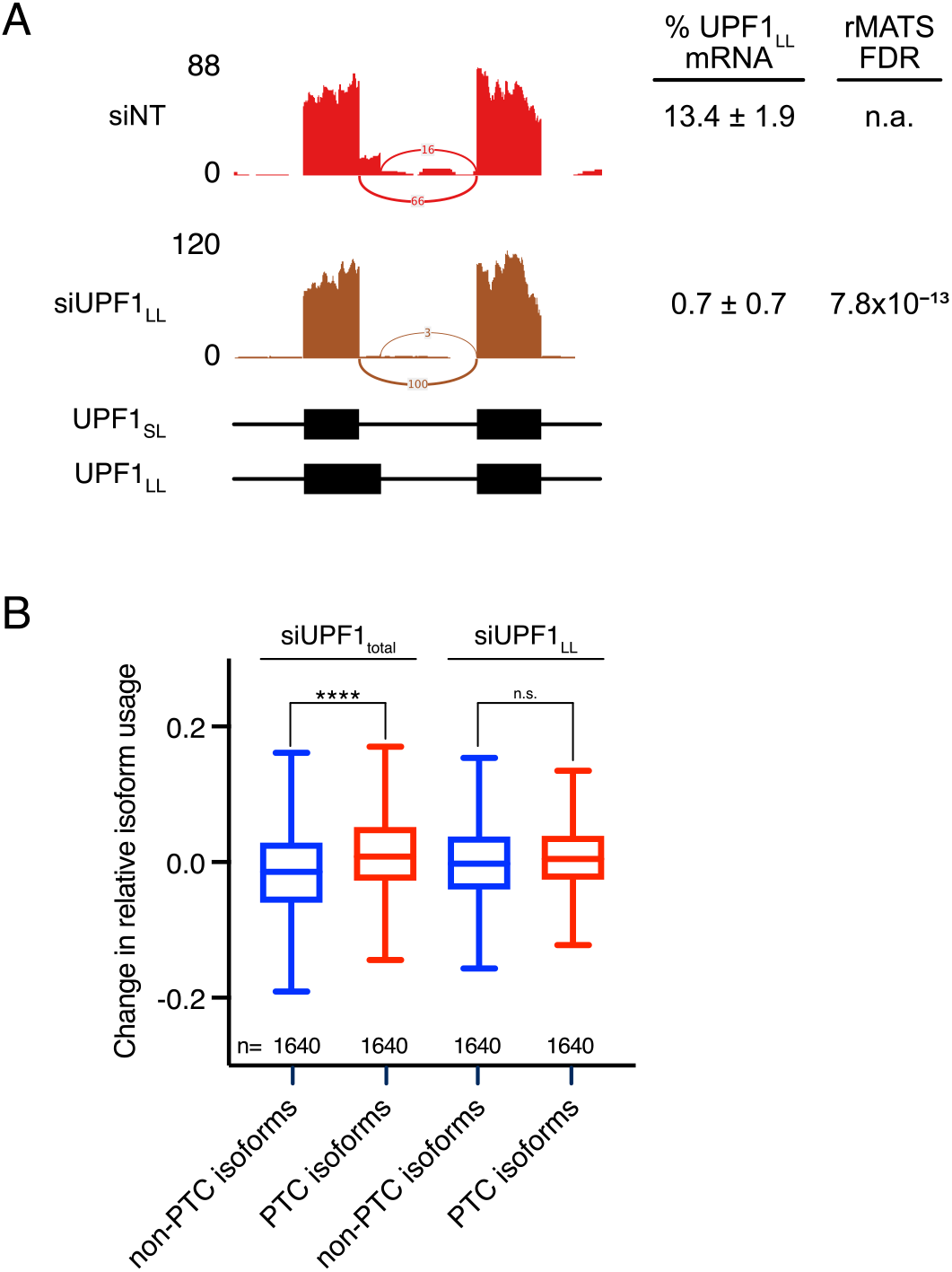
Isoform-specific depletion of UPF1_LL_ does not systematically affect EJC-enhanced NMD. **A.** Sashimi plot from representative RNA-seq samples of siNT and siUPF1_LL_ knockdown cells. Percent spliced in values and FDR were calculated with rMATS software. **B.** Box plot of the change in relative isoform usage of PTC-containing transcripts as determined from RNA-seq following total UPF1 (siUPF_total_) or UPF1_LL_-specific knockdown in HEK-293 cells. Genes for which expression of an isoform containing a termination codon greater than 50 nt upstream of the final exon junction wereanalyzed, with mRNA isoforms classified as having or lacking a premature termination codon (PTC). Statistical significance was determined by one-way ANOVA (**** *P<* 1×10^−15^). Boxes indicate interquartileranges, and bars indicate Tukey whiskers.

**Figure S2.**
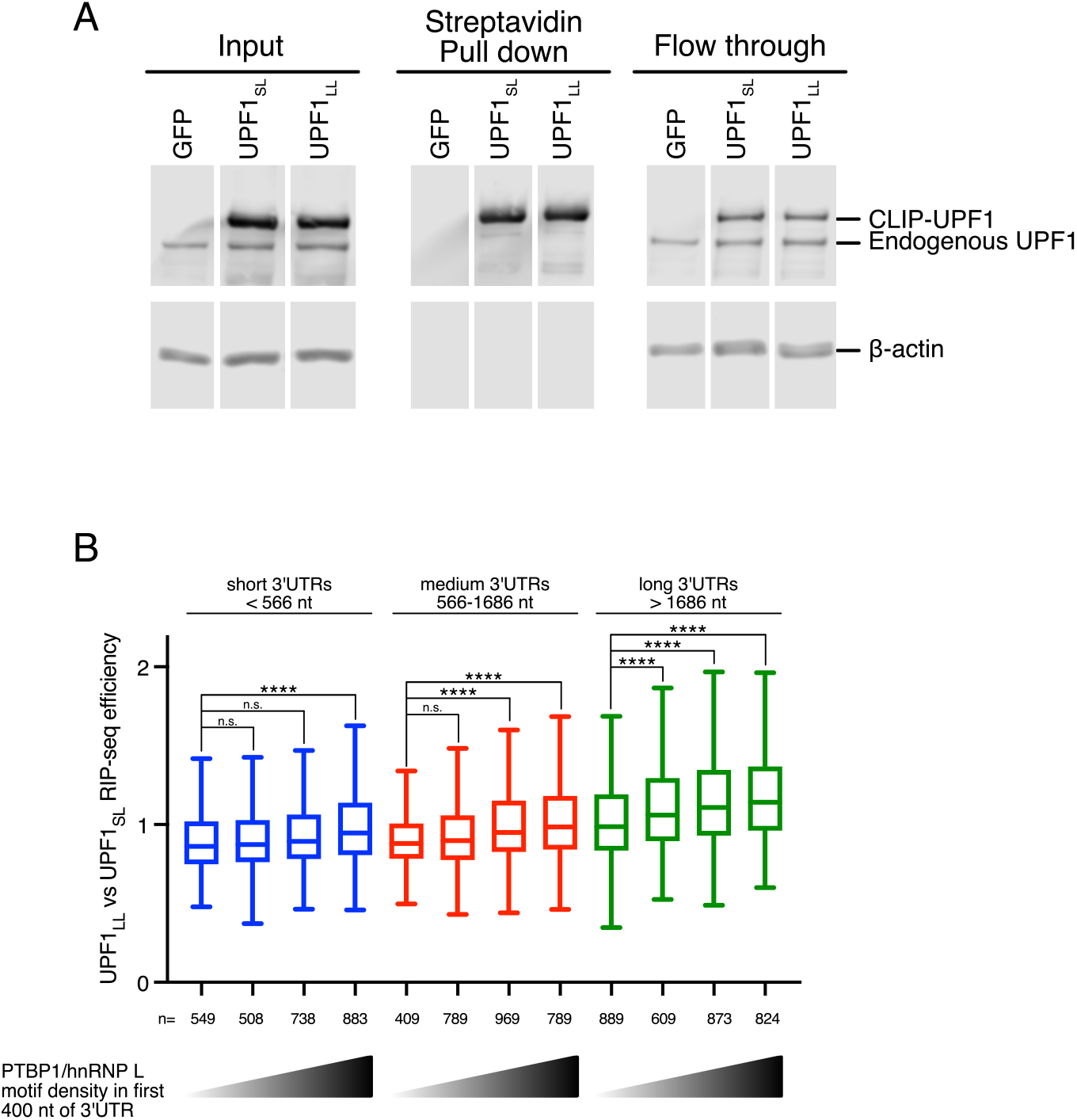
UPF1_LL_ can bind transcripts normally shielded by protective proteins. **A.** Western blots of CLIP-UPF1_SL_ and CLIP-UPF1_LL_ cell lysates before (input) and after (pull down) biotin-streptavidin affinity purification. Input and flow through lanes represent 5% of total material. **B.** Box plot of recovered mRNAs as determined from RIP-seq efficiency in CLIP-UPF1_LL_ vs CLIP- UPF1_SL_ affinity purifications. mRNAs were binned according to 3’UTR length (short, medium, long) and then equally subdivided by PTBP1 and/or hnRNP L motif density within the first 400 nt of 3’UTR, as indicated by the gradient triangles. Statistical significance was determined by K-W test, with Dunn’s correction for multiple comparisons (**** *P* < 6×10^−5^). Boxes indicate interquartile ranges, and bars indicate Tukey whiskers.

**Figure S3.**
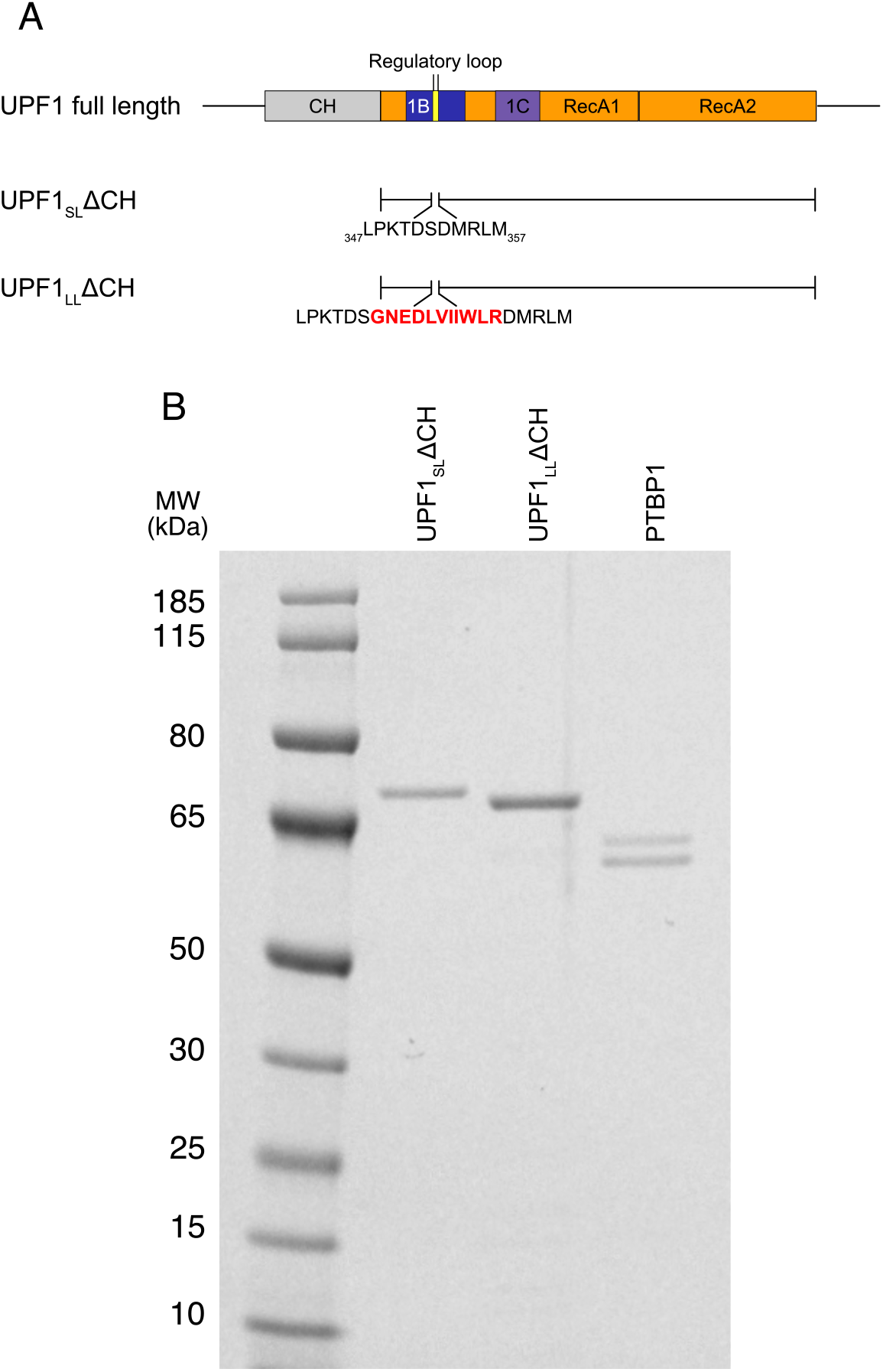
Reduced sensitivity of UPF1_LL_ to translocation inhibition by PTBP1. **A.** Schematic representation of the domain organization of full-length human UPF1, with the UPF1ΔCH constructs assessed in this study depicted below. The 11 amino acid extension in the regulatory loop of UPF1_LL_ is indicated in red text. **B.** SDS-PAGE of recombinant 6xH1S-CBP-UPF1_SL_ΔCH (MW: 76 kDa), 6xH1S-UPF1_LL_ΔCH (MW: 73 kDa), and 6xH1S-PTBP1 (MW: 62 kDa) proteins purified from *E.coli* and used in this study. PTBP1 doublet consists of differentially migrating conformers of identical MW, as determined by mass spectrometry of bands excised from SDS-PAGE gels (Fritz et al., 2020).

**Figure S4.**
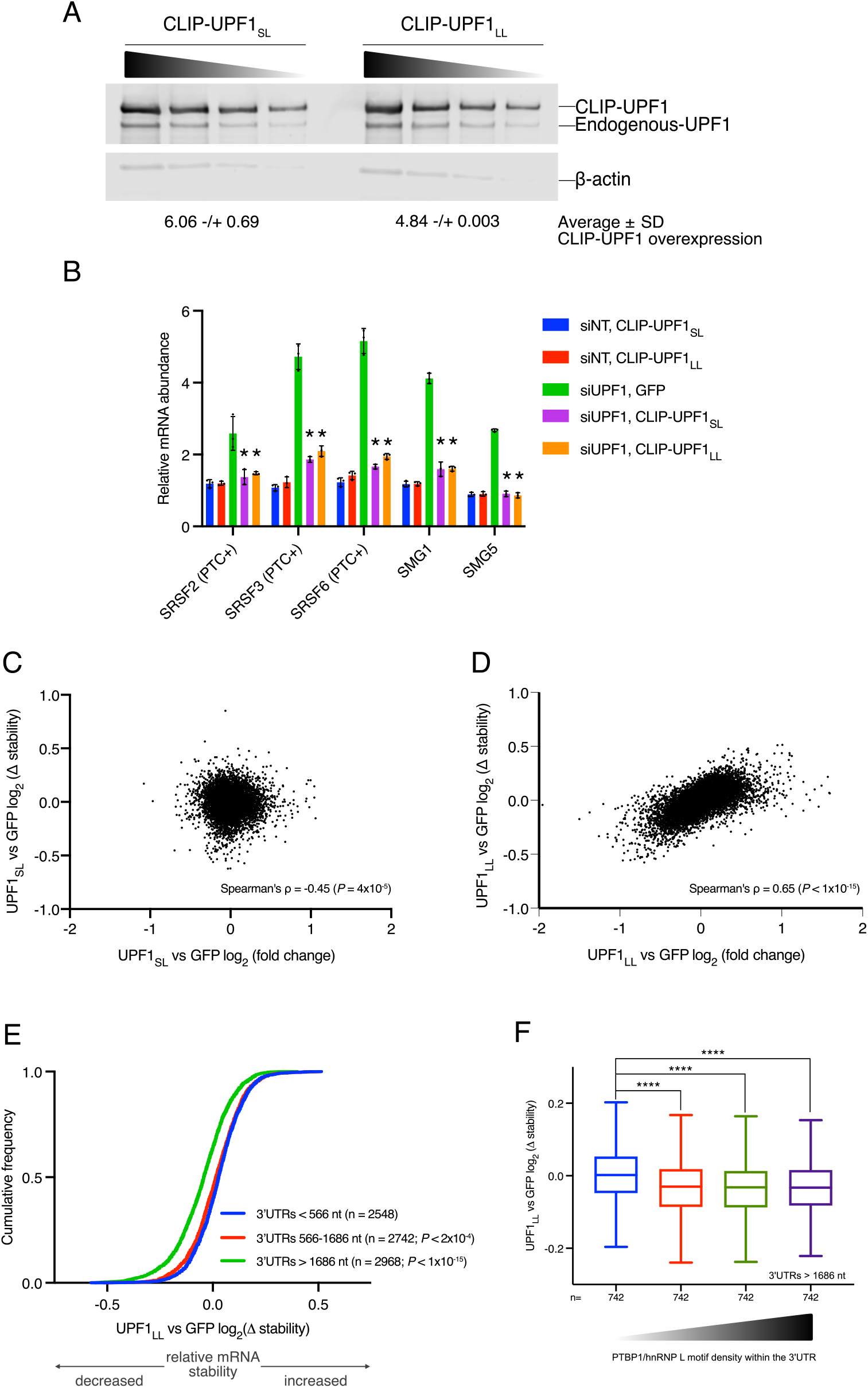
NMD protection can be overcome by UPF1_LL_. **A.** Western blots of CLIP-UPF1_SL_ and CLIP-UPF1_LL_ overexpression. Membranes were probed with an anti-UPF1 antibody that detects both endogenous and CLIP-tagged UPF1. Wedge indicates serial two- fold dilutions of lysate. Average (± standard deviation) of CLIP-UPF1 overexpression was determined from two replicate membranes. **B.** RT-qPCR analysis of well-characterized NMD targets following total UPF1 knockdown and rescue with siRNA-resistant CLIP-tagged UPF1. Relative fold changes are in reference to the GFP-expressing control line treated with a non-targeting (NT) siRNA. Significance of NMD rescue by CLIP-UPF1 was compared to the GFP-expressing control line treated with total UPF1 siRNA. Asterisk(*) indicates *P* < 0.0001, as determined by two-way ANOVA with multiple comparisons. Error bars indicate SD (n = 3). PTC+ indicates the use of primers specific to transcript isoforms with validated poison exons (Lareau et al., 2007; Ni et al., 2007). **C.** Scatterplot of changes in relative mRNA stability as determined by REMBRANDTS analysis of RNA- seq vs relative mRNA abundance as determined by kallisto, comparing CLIP-UPF1_SL_ to GFP overexpression. **D.** Scatterplot as in (C), comparing CLIP-UPF1_LL_ to GFP overexpression. **E.** CDF plot of relative mRNA stability as determined by REMBRANDTS analysis of RNA-seq following CLIP-UPF1_LL_ or GFP overexpression. mRNAs were binned according to 3’UTR lengths (short, medium, long). Statistical significance was determined by K-W test, with Dunn’s correction for multiple comparisons. **F.** Box plot of relative mRNA stability as determined by REMBRANDTS analysis of RNA-seq for transcripts having long 3’UTRs (> 1578 nt) following CLIP-UPF1_LL_ or GFP overexpression. mRNAs were binned by PTBP1 and/or hnRNP L motif density within the 3’UTR. Statistical significance was determined by K-W test, with Dunn’s correction for multiple comparisons (**** *P* < 3×10^−14^). Boxes indicate interquartile ranges, and bars indicate Tukey whiskers.

**Figure S5.**
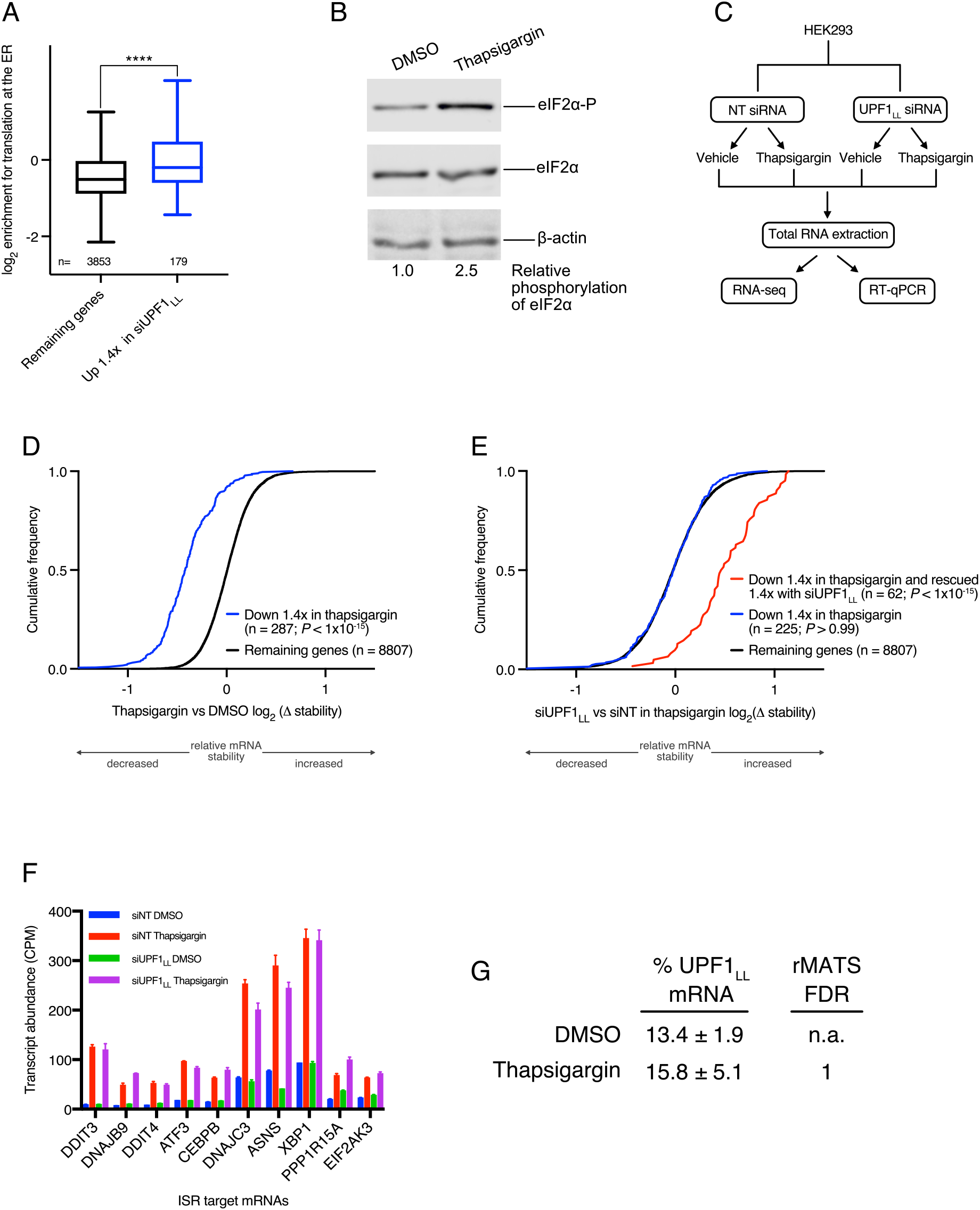
Transcripts targeted by UPF1_LL_ are coordinately down-regulated during ER stress and induction of the ISR. **A.** Box plot of log_2_ enrichment for translation at the ER (Jan et al., 2014). mRNAs were binned by sensitivity to UPF1_LL_-specific knockdown in HEK-293 cells. Statistical significance was determined by K- S test (**** *P* = 1×10^−6^). Boxes indicate interquartile ranges, and bars indicate Tukey whiskers. **B.** Western blot of elF2a phosphorylation following treatment of HEK-293 cells with 1 *μ*M thapsigargin for 6hr. **C.** Schematic of the RNA-seq experimental workflow and conditions for UPF1_LL_ knockdown and thapsigargin treatment. **D.** CDF plot of relative mRNA stability as determined by REMBRANDTS analysis of RNA-seq following treatment of HEK-293 cells with 1 *μ*M thapsigargin for 6hr. mRNAs were binned by changes in relative mRNA abundance in thapsigargin. Statistical significance was determined by K-S test. **E.** CDF plot of relative mRNA stability as determined by REMBRANDTS analysis of RNA-seq following UPF1_LL_ knockdown in HEK-293 cells and treatment with 1 *μ*M thapsigargin for 6hr. mRNAs were binned by changes in relative mRNA abundance in thapsigargin with UPF1_LL_ knockdown. Statistical significance was determined by K-W test, with Dunn’s correction for multiple comparisons. **F.** Quantification of characterized ISR-target transcript abundance in RNA-seq of the indicated conditions. Error bars indicate SD (n = 3). **G.** Quantification of UPF1_LL_ isoform expression in control and thapsigargin-treated HEK-293 cells from rMATS analyses.

**Figure S6.**
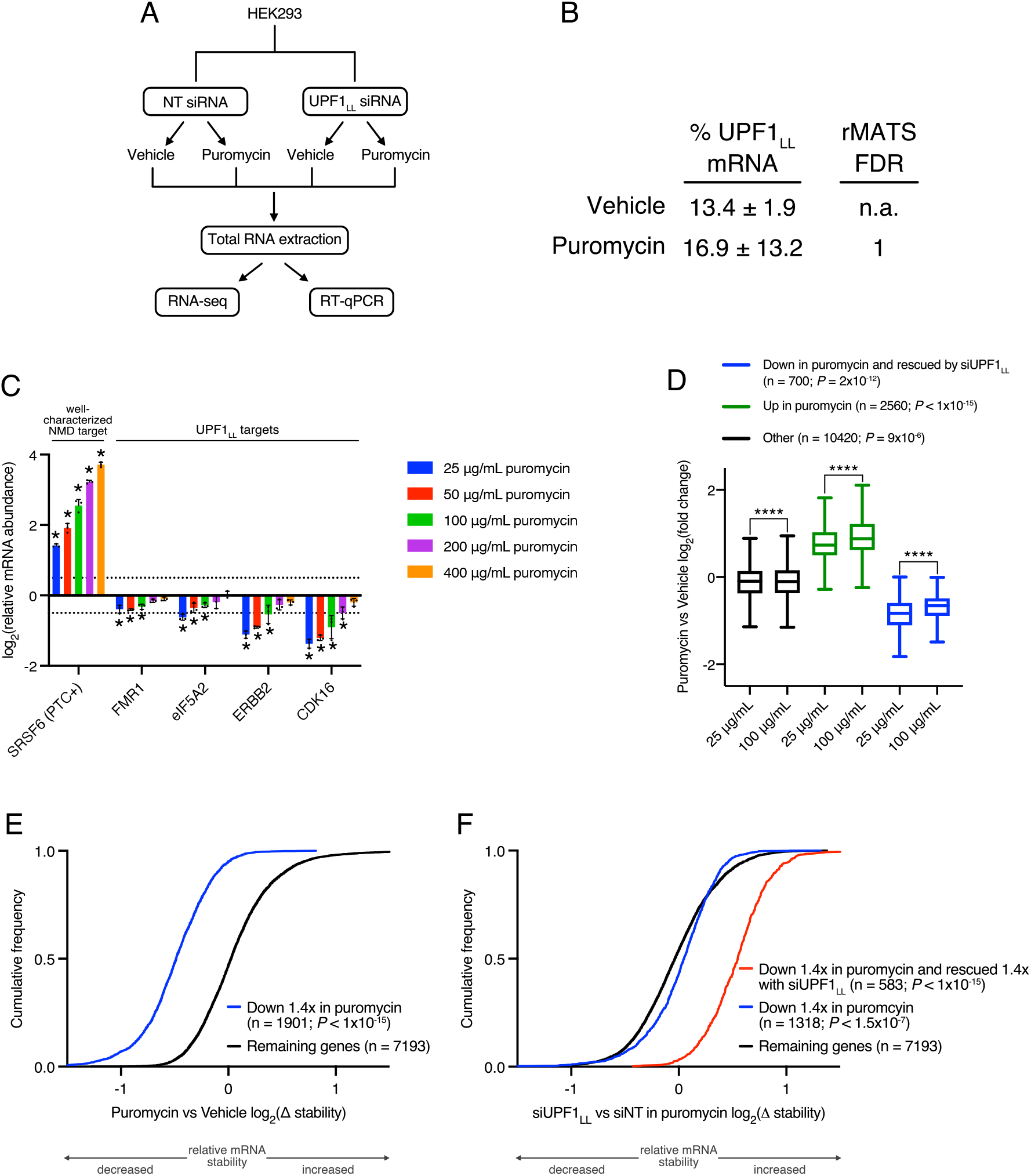
Decreased translation promotes UPF1_LL_ activity. **A.** Schematic of the RNA-seq experimental workflow and conditions for UPF1_LL_ knockdown and puromycin treatment. **B.** Quantification of UPF1_LL_ isoform expression in control and puromycin-treated HEK-293 cells from rMATS analyses. **C.** RT-qPCR analysis of indicated transcripts following treatment of HEK-293 cells with indicated concentrations of puromycin for 4hr. Relative fold changes are in reference to vehicle-treated control. Significance of puromycin treatment on relative transcript abundance was compared to the vehicle- treated control. Asterisk (*) indicates *P* < 0.05, as determined by two-way ANOVA. Error bars indicate SD (n = 3). Dashed lines indicate log_2_(fold change) of ± 0.5. PTC+ indicates the use of primers specific to the transcript isoform with a validated poison exon (Lareau et al., 2007; Ni et al., 2007). **D.** Box plot of relative mRNA abundance as determined by RNA-seq following treatment of HEK-293 cells with 25 μg/ml or 100 μg/ml puromycin. mRNAs were binned according to sensitivity to 50 μg/ml puromycin and UPF1_LL_ knockdown. Statistical significance was determined by K-S test (**** *P* < 9×10^−6^). Boxes indicate interquartile ranges, and bars indicate Tukey whiskers. **E.** CDF plot of relative mRNA stability as determined by REMBRANDTS analysis of RNA-seq following treatment of HEK-293 cells with 50 *μ*g/ml puromycin for 4hr. mRNAs were binned by changes in relative mRNA abundance in puromycin. Statistical significance was determined by K-S test. **F.** CDF plot of relative mRNA stability as determined by REMBRANDTS analysis of RNA-seq following UPF1_LL_ knockdown in HEK-293 cells and treatment with 50 μg/ml puromycin for 4hr. mRNAs were binned by changes in relative mRNA abundance in puromycin with UPF1_LL_ knockdown. Statistical significance was determined by K-S test, with Dunn’s correction for multiple comparisons.

## References

Akopian D, Shen K, Zhang X & Shan S-O (2013) Signal recognition particle: an essential protein-targeting machine. Annu Rev Biochem 82: 693–721

Alkallas R, Fish L, Goodarzi H & Najafabadi HS (2017) Inference of RNA decay rate from transcriptional profiling highlights the regulatory programs of Alzheimer’s disease. Nat Commun 8: 909

Anders S, Reyes A & Huber W (2012) Detecting differential usage of exons from RNA-seq data. Genome Res 22: 2008–2017

Ashburner M, Ball CA, Blake JA, Botstein D, Butler H, Cherry JM, Davis AP, Dolinski K, Dwight SS, Eppig JT, et al (2000) Gene ontology: tool for the unification of biology. The Gene Ontology Consortium. Nat Genet 25: 25–29

Baird TD, Cheng KC-C, Chen Y-C, Buehler E, Martin SE, Inglese J & Hogg JR (2018) ICE1 promotes the link between splicing and nonsense-mediated mRNA decay. Elife 7: e33178

Baird TD & Wek RC (2012) Eukaryotic initiation factor 2 phosphorylation and translational control in metabolism. Adv Nutr 3: 307–321

Baker SL & Hogg JR (2017) A system for coordinated analysis of translational readthrough and nonsense-mediated mRNA decay. PLoS One 12: e0173980

Bray NL, Pimentel H, Melsted P & Pachter L (2016) Near-optimal probabilistic RNA-seq quantification. Nat Biotechnol 34: 525–527

Carter MS, Doskow J, Morris P, Li S, Nhim RP, Sandstedt S & Wilkinson MF (1995) A regulatory mechanism that detects premature nonsense codons in T-cell receptor transcripts in vivo is reversed by protein synthesis inhibitors in vitro. J Biol Chem 270: 28995–29003

Causier B, Li Z, De Smet R, Lloyd JPB, Van de Peer Y & Davies B (2017) Conservation of Nonsense-Mediated mRNA Decay Complex Components Throughout Eukaryotic Evolution. Sci Rep 7: 16692

Chakrabarti S, Jayachandran U, Bonneau F, Fiorini F, Basquin C, Domcke S, Le Hir H & Conti E (2011) Molecular mechanisms for the RNA-dependent ATPase activity of Upf1 and its regulation by Upf2. Mol Cell 41: 693–703

Chamieh H, Ballut L, Bonneau F & Le Hir H (2008) NMD factors UPF2 and UPF3 bridge UPF1 to the exon junction complex and stimulate its RNA helicase activity. Nat Struct Mol Biol 15: 85–93

Chan WK, Huang L, Gudikote JP & Chang YF (2007) An alternative branch of the nonsense‐mediated decay pathway. EMBO J

Cheng Z, Muhlrad D, Lim MK, Parker R & Song H (2007) Structural and functional insights into the human Upf1 helicase core. EMBO J 26: 253–264

Chen S, Zhou Y, Chen Y & Gu J (2018) fastp: an ultra-fast all-in-one FASTQ preprocessor. Bioinformatics 34: i884–i890

Colombo M, Karousis ED, Bourquin J, Bruggmann R & Mühlemann O (2017) Transcriptome-wide identification of NMD-targeted human mRNAs reveals extensive redundancy between SMG6- and SMG7-mediated degradation pathways. RNA 23: 189–201 doi:10.1261/rna.059055.116 [PREPRINT]

Costa-Mattioli M & Walter P (2020) The integrated stress response: From mechanism to disease. Science 368

Czaplinski K, Weng Y, Hagan KW & Peltz SW (1995) Purification and characterization of the Upf1 protein: a factor involved in translation and mRNA degradation. RNA 1: 610–623

Dockendorff TC & Labrador M (2019) The Fragile X Protein and Genome Function. Mol Neurobiol 56: 711–721

Durand S, Franks TM & Lykke-Andersen J (2016) Hyperphosphorylation amplifies UPF1 activity to resolve stalls in nonsense-mediated mRNA decay. Nat Commun 7: 12434

Eberle AB, Lykke-Andersen S, Mühlemann O & Jensen TH (2009) SMG6 promotes endonucleolytic cleavage of nonsense mRNA in human cells. Nat Struct Mol Biol 16: 49–55

Eden E, Navon R, Steinfeld I, Lipson D & Yakhini Z (2009) GOrilla: a tool for discovery and visualization of enriched GO terms in ranked gene lists. BMC Bioinformatics 10: 48

Fiorini F, Bonneau F & Le Hir H (2012) Biochemical characterization of the RNA helicase UPF1 involved in nonsense-mediated mRNA decay. Methods Enzymol 511: 255–274

Fiorini F, Boudvillain M & Le Hir H (2013) Tight intramolecular regulation of the human Upf1 helicase by its N- and C-terminal domains. Nucleic Acids Res 41: 2404–2415

Franks TM, Singh G & Lykke-Andersen J (2010) Upf1 ATPase-dependent mRNP disassembly is required for completion of nonsense- mediated mRNA decay. Cell 143: 938–950

Fritz SE, Haque N & Hogg JR (2018) Highly efficient in vitro translation of authentic affinity-purified messenger ribonucleoprotein complexes. RNA 24: 982–989

Fritz SE, Ranganathan S, Wang CD & Hogg JR (2020) The RNA-binding protein PTBP1 promotes ATPase-dependent dissociation of the RNA helicase UPF1 to protect transcripts from nonsense-mediated mRNA decay. J Biol Chem 295: 11613–11625

Gautier A, Juillerat A, Heinis C, Corrêa IR Jr, Kindermann M, Beaufils F & Johnsson K (2008) An engineered protein tag for multiprotein labeling in living cells. Chem Biol 15: 128–136

Ge Z, Quek BL, Beemon KL & Robert Hogg J (2016) Polypyrimidine tract binding protein 1 protects mRNAs from recognition by the nonsense-mediated mRNA decay pathway. eLife Sciences 5: e11155

Goetz AE & Wilkinson M (2017) Stress and the nonsense-mediated RNA decay pathway. Cell Mol Life Sci 74: 3509–3531

Goldstein JL & Brown MS (2009) The LDL receptor. Arterioscler Thromb Vasc Biol 29: 431–438

Gowravaram M, Bonneau F, Kanaan J, Maciej VD, Fiorini F, Raj S, Croquette V, Le Hir H & Chakrabarti S (2018) A conserved structural element in the RNA helicase UPF1 regulates its catalytic activity in an isoform-specific manner. Nucleic Acids Res 46: 2648–2659

Harbeck N, Penault-Llorca F, Cortes J, Gnant M, Houssami N, Poortmans P, Ruddy K, Tsang J & Cardoso F (2019) Breast cancer. Nat Rev Dis Primers 5: 66

Heinz S, Benner C, Spann N, Bertolino E, Lin YC, Laslo P, Cheng JX, Murre C, Singh H & Glass CK (2010) Simple combinations of lineage-determining transcription factors prime cis-regulatory elements required for macrophage and B cell identities. Mol Cell 38: 576–589

Hodgkin J, Papp A, Pulak R, Ambros V & Anderson P (1989) A new kind of informational suppression in the nematode Caenorhabditis elegans. Genetics 123: 301–313

Hogg JR & Collins K (2007a) RNA-based affinity purification reveals 7SK RNPs with distinct composition and regulation. RNA 13: 868–880

Hogg JR & Collins K (2007b) Human Y5 RNA specializes a Ro ribonucleoprotein for 5S ribosomal RNA quality control. Genes Dev 21: 3067–3072

Hogg JR & Goff SP (2010) Upf1 senses 3′ UTR length to potentiate mRNA decay. Cell 143: 379–389

Hollien J & Weissman JS (2006) Decay of endoplasmic reticulum-localized mRNAs during the unfolded protein response. Science 313: 104–107

Huang L, Lou C-H, Chan W, Shum EY, Shao A, Stone E, Karam R, Song H-W & Wilkinson MF (2011) RNA homeostasis governed by cell type-specific and branched feedback loops acting on NMD. Mol Cell 43: 950–961

Huntzinger E, Kashima I, Fauser M, Saulière J & Izaurralde E (2008) SMG6 is the catalytic endonuclease that cleaves mRNAs containing nonsense codons in metazoan. RNA 14: 2609–2617

Hurt JA, Robertson AD & Burge CB (2013) Global analyses of UPF1 binding and function reveal expanded scope of nonsense-mediated mRNA decay. Genome Res 23: 1636–1650

Jan CH, Williams CC & Weissman JS (2014) Principles of ER cotranslational translocation revealed by proximity-specific ribosome profiling. Science 346

Karousis ED & Mühlemann O (2018) Nonsense-Mediated mRNA Decay Begins Where Translation Ends. Cold Spring Harb Perspect Biol

Kashima I, Yamashita A, Izumi N, Kataoka N, Morishita R, Hoshino S, Ohno M, Dreyfuss G & Ohno S (2006) Binding of a novel SMG-1-Upf1-eRF1-eRF3 complex (SURF) to the exon junction complex triggers Upf1 phosphorylation and nonsense-mediated mRNA decay. Genes Dev 20: 355–367

Keene JD (2007) RNA regulons: coordination of post-transcriptional events. Nat Rev Genet 8: 533–543

Kim D, Paggi JM, Park C, Bennett C & Salzberg SL (2019) Graph-based genome alignment and genotyping with HISAT2 and HISAT-genotype. Nat Biotechnol 37: 907–915

Kim YK & Maquat LE (2019) UPFront and center in RNA decay: UPF1 in nonsense-mediated mRNA decay and beyond. RNA 25: 407–422

Kishor A, Fritz SE, Haque N, Ge Z, Tunc I, Yang W, Zhu J & Hogg JR (2020) Activation and inhibition of nonsense-mediated mRNA decay control the abundance of alternative polyadenylation products. Nucleic Acids Res 48: 7468–7482

Kishor A, Fritz SE & Robert Hogg J (2019) Nonsense‐mediated mRNA decay: The challenge of telling right from wrong in a complex transcriptome. Wiley Interdisciplinary Reviews: RNA 10 doi:10.1002/wrna.1548 [PREPRINT]

Kishor A, Ge Z & Hogg JR (2018) hnRNP L-dependent protection of normal mRNAs from NMD subverts quality control in B cell lymphoma. EMBO J

Kurosaki T, Li W, Hoque M, Popp MW-L, Ermolenko DN, Tian B & Maquat LE (2014) A post-translational regulatory switch on UPF1 controls targeted mRNA degradation. Genes Dev 28: 1900–1916

Lareau LF, Inada M, Green RE, Wengrod JC & Brenner SE (2007) Unproductive splicing of SR genes associated with highly conserved and ultraconserved DNA elements. Nature 446: 926–929

Lavysh D & Neu-Yilik G (2020) UPF1-Mediated RNA Decay-Danse Macabre in a Cloud. Biomolecules 10: 999

Lee SR, Pratt GA, Martinez FJ, Yeo GW & Lykke-Andersen J (2015) Target Discrimination in Nonsense-Mediated mRNA Decay Requires Upf1 ATPase Activity. Mol Cell 59: 413–425

Le Hir H, Izaurralde E, Maquat LE & Moore MJ (2000) The spliceosome deposits multiple proteins 20-24 nucleotides upstream of mRNA exon-exon junctions. EMBO J 19: 6860–6869

Le Hir H, Moore MJ & Maquat LEM (2000) Pre-mRNA splicing alters mRNP composition: evidence for stable association of proteins at exon-exon junctions. Genes Dev 14: 1098–1108

Liao Y, Smyth GK & Shi W (2014) featureCounts: an efficient general purpose program for assigning sequence reads to genomic features. Bioinformatics 30: 923–930

Li Z, Vuong JK, Zhang M, Stork C & Zheng S (2017) Inhibition of nonsense-mediated RNA decay by ER stress. RNA 23: 378–394

Loh B, Jonas S & Izaurralde E (2013) The SMG5-SMG7 heterodimer directly recruits the CCR4-NOT deadenylase complex to mRNAs containing nonsense codons via interaction with POP2. Genes Dev 27: 2125–2138

Longman D, Jackson-Jones KA, Maslon MM, Murphy LC, Young RS, Stoddart JJ, Hug N, Taylor MS, Papadopoulos DK & Cáceres JF (2020) Identification of a localized nonsense-mediated decay pathway at the endoplasmic reticulum. Genes Dev

Martinez-Nunez RT, Wallace A, Coyne D, Jansson L, Rush M, Ennajdaoui H, Katzman S, Bailey J, Deinhardt K, Sanchez-Elsner T, et al (2017) Modulation of nonsense mediated decay by rapamycin. Nucleic Acids Res 45: 3448–3459

Martin M (2011) Cutadapt removes adapter sequences from high-throughput sequencing reads. EMBnet.journal 17: 10–12

Mathews MB & Hershey JWB (2015) The translation factor eIF5A and human cancer. Biochim Biophys Acta 1849: 836–844

Michlewski G & Cáceres JF (2019) Post-transcriptional control of miRNA biogenesis. RNA 25: 1–16

Mugridge JS, Coller J & Gross JD (2018) Structural and molecular mechanisms for the control of eukaryotic 5’-3’ mRNA decay. Nat Struct Mol Biol 25: 1077–1085

Nathans D (1964) Puromycin inhibition of protein synthesis: incorporation of puromycin into peptide chains. Proc Natl Acad Sci U S A 51: 585–592

Nicholson P, Josi C, Kurosawa H, Yamashita A & Mühlemann O (2014) A novel phosphorylation-independent interaction between SMG6 and UPF1 is essential for human NMD. Nucleic Acids Res 42: 9217–9235

Nickless A, Jackson E, Marasa J, Nugent P, Mercer RW, Piwnica-Worms D & You Z (2014) Intracellular calcium regulates nonsense-mediated mRNA decay. Nat Med 20: 961–966

Ni JZ, Grate L, Donohue JP, Preston C, Nobida N, O’Brien G, Shiue L, Clark TA, Blume JE & Ares M Jr (2007) Ultraconserved elements are associated with homeostatic control of splicing regulators by alternative splicing and nonsense-mediated decay. Genes Dev 21: 708–718

Niknafs YS, Pandian B, Iyer HK, Chinnaiyan AM & Iyer MK (2017) TACO produces robust multisample transcriptome assemblies from RNA-seq. Nat Methods 14: 68–70

Page MFP, Carr BC, Anders KRA, Grimson AG & Anderson PA (1999) SMG-2 is a phosphorylated protein required for mRNA surveillance in Caenorhabditis elegans and related to Upf1p of yeast. Mol Cell Biol 19: 5943–5951

Pakos-Zebrucka K, Koryga I, Mnich K, Ljujic M, Samali A & Gorman AM (2016) The integrated stress response. EMBO Rep 17: 1374–1395

Paz I, Kosti I, Ares M Jr, Cline M & Mandel-Gutfreund Y (2014) RBPmap: a web server for mapping binding sites of RNA-binding proteins. Nucleic Acids Res 42: W361–7

Pertea M, Pertea GM, Antonescu CM, Chang T-C, Mendell JT & Salzberg SL (2015) StringTie enables improved reconstruction of a transcriptome from RNA-seq reads. Nat Biotechnol 33: 290–295

Powell D (2015) Degust: Visualize, explore and appreciate RNA-seq differential gene-expression data. In COMBINE RNA-seq workshop

Pulak R & Anderson P (1993) mRNA surveillance by the Caenorhabditis elegans smg genes. Genes Dev 7: 1885–1897

Risso D, Ngai J, Speed TP & Dudoit S (2014) Normalization of RNA-seq data using factor analysis of control genes or samples. Nat Biotechnol 32: 896–902

Ritchie ME, Phipson B, Wu D, Hu Y, Law CW, Shi W & Smyth GK (2015) limma powers differential expression analyses for RNA-sequencing and microarray studies. Nucleic Acids Res 43: e47

Robinson MD, McCarthy DJ & Smyth GK (2009) edgeR : a Bioconductor package for differential expression analysis of digital gene expression data. Bioinformatics 26: 139–140

Schofield JA, Duffy EE, Kiefer L, Sullivan MC & Simon MD (2018) TimeLapse-seq: adding a temporal dimension to RNA sequencing through nucleoside recoding. Nat Methods 15: 221–225

Shen S, Park JW, Lu Z-X, Lin L, Henry MD, Wu YN, Zhou Q & Xing Y (2014) rMATS: robust and flexible detection of differential alternative splicing from replicate RNA-Seq data. Proc Natl Acad Sci U S A 111: E5593–601

Singh G, Rebbapragada I & Lykke-Andersen J (2008) A competition between stimulators and antagonists of Upf complex recruitment governs human nonsense-mediated mRNA decay. PLoS Biol 6: e111

Smith JE & Baker KE (2015) Nonsense-mediated RNA decay - a switch and dial for regulating gene expression. Bioessays 37: 612–623

Song MS, Salmena L & Pandolfi PP (2012) The functions and regulation of the PTEN tumour suppressor. Nat Rev Mol Cell Biol 13: 283–296

The Gene Ontology Consortium (2019) The Gene Ontology Resource: 20 years and still GOing strong. Nucleic Acids Res 47: D330–D338

Thorvaldsdóttir H, Robinson JT & Mesirov JP (2013) Integrative Genomics Viewer (IGV): high-performance genomics data visualization and exploration. Brief Bioinform 14: 178–192

Toma KG, Rebbapragada I, Durand S & Lykke-Andersen J (2015) Identification of elements in human long 3’ UTRs that inhibit nonsense-mediated decay. RNA 21: 887–897

Vitting-Seerup K & Sandelin A (2019) IsoformSwitchAnalyzeR: Analysis of changes in genome-wide patterns of alternative splicing and its functional consequences. Bioinformatics: ii: btz247

Watson M, Park Y & Thoreen C (2020) Roadblock-qPCR: A simple and inexpensive strategy for targeted measurements of mRNA stability. RNA

Wek RC (2018) Role of eIF2α Kinases in Translational Control and Adaptation to Cellular Stress. Cold Spring Harb Perspect Biol 10

Weng Y, Czaplinski K & Peltz SW (1998) ATP is a cofactor of the Upf1 protein that modulates its translation termination and RNA binding activities. RNA 4: 205–214

Yepiskoposyan H, Aeschimann F, Nilsson D, Okoniewski M & Mühlemann O (2011) Autoregulation of the nonsense-mediated mRNA decay pathway in human cells. RNA 17: 2108–2118

Yi Z, Sanjeev M & Singh G (2020) The Branched Nature of the Nonsense-Mediated mRNA Decay Pathway. Trends Genet

Young-Baird SK, Lourenço MB, Elder MK, Klann E, Liebau S & Dever TE (2020) Suppression of MEHMO Syndrome Mutation in eIF2 by Small Molecule ISRIB. Mol Cell 77: 875–886.e7

Young SK & Wek RC (2016) Upstream Open Reading Frames Differentially Regulate Gene-specific Translation in the Integrated Stress Response. J Biol Chem 291: 16927–16935

Zünd D, Gruber AR, Zavolan M & Mühlemann O (2013) Translation-dependent displacement of UPF1 from coding sequences causes its enrichment in 3’ UTRs. Nat Struct Mol Biol 20: 936–943

